# Anisomycin-induced ribosome collisions follow a dose-dependent positional rule, distinct from the UV-induced collision landscape

**DOI:** 10.64898/2026.10.12.758323

**Authors:** Sébastien Chamois, Julien Soudet, Virginie Ricci, Félix Naef, Cédric Gobet, David Gatfield

**Author notes:** corresponding author e-mail addresses.

## Abstract

Ribosome collisions trigger stress pathways such as the ribotoxic stress response, yet how different ribosome-impairing stimuli shape where collisions form on mRNAs remains unclear. Here, we combine monosome- and disome-resolved ribosome profiling with proteomics of collided-ribosome fractions to compare anisomycin- and UV-induced ribosome impairment in mouse embryonic stem cells. Low-dose anisomycin, a widely used collision-inducing paradigm, does not generate collisions dispersed along transcripts but a post-initiation collision zone, with stalled monosomes and disomes accumulating within the first ∼70 nucleotides of coding sequences across most translated mRNAs. High-dose anisomycin, commonly considered collision-poor, produces the same pattern shifted closer to the start codon and reduced in amplitude. These observations support a single dose-dependent positional rule, in which drug occupancy determines how far ribosomes progress before arrest and ongoing initiation supplies trailing ribosomes. In contrast, 30 min after UV irradiation, collisions are detectable biochemically but appear more dispersed, with few recurrent hotspots and preferential accumulation on highly translated mRNAs. Collided-ribosome fractions share enrichment of the collision sensor EDF1 across conditions, whereas recruitment of ZNF598 and initiation factors is most evident after UV. Our results refine the interpretation of anisomycin-based collision paradigms and show that collision landscapes depend on the nature of the stimulus.

## Introduction

Beyond its core function in protein synthesis (1), the ribosome has emerged as a central signalling platform that links translational activity to stress-response pathways. Perturbations of translation engage the integrated stress response (ISR), which attenuates translation initiation through phosphorylation of eIF2α (2); ribosome-associated quality control (RQC), which resolves stalled translation complexes and degrades aberrant nascent peptides (3); and the ribotoxic stress response (RSR), which activates MAP kinase cascades that can induce cell-cycle arrest and cell death (4).

Ribosome collisions have emerged as key molecular signals connecting translational perturbations to these pathways (5). Collisions occur transiently even during unperturbed translation elongation (6), but persistent slowing or stalling causes collided ribosomes to accumulate, and these are recognised by dedicated sensors. Collided ribosomes activate the ribosome-associated MAP3 kinase ZAKα, which initiates p38 and JNK signalling (7,8), and they serve as initiating signals for the recruitment of RQC and other surveillance factors.

Much of our understanding of collision-induced signalling derives from translation elongation inhibitors such as anisomycin (ANS) or emetine (9,10). The prevailing model posits that low inhibitor concentrations stall a sparse subset of ribosomes dispersed along transcripts, so that upstream ribosomes continue to elongate and collide with them, whereas high concentrations immobilise most elongating ribosomes simultaneously and largely prevent collisions. This model makes clear predictions: collisions under low-dose ANS should be broadly distributed along coding sequences, and high-dose ANS should be essentially collision-free. Yet, despite the widespread use of this paradigm, these predictions have not been tested transcriptome-wide in mammalian cells, and it remains unknown where along coding sequences, and on which mRNAs, drug-induced collisions actually form.

A related question is whether collision landscapes induced by elongation inhibitors resemble those arising from other forms of translational stress. Ultraviolet B (UVB) irradiation provides an informative comparison: it impairs translation through RNA photolesions rather than by direct inhibition of the ribosome (11), activates the RSR and produces biochemical signatures of ribosome collisions (12,13), and additionally engages the ISR (14). How UV-induced collisions are distributed across the translatome, however, remains poorly defined.

Here, we combine monosome- and disome-resolved ribosome profiling (Ribo-seq) with proteomics of collided-ribosome fractions in mouse embryonic stem cells (mESCs) to compare low- and high-dose ANS with UV irradiation. We find that low-dose ANS does not produce dispersed collisions but a post-initiation collision zone within the first codons of most translated mRNAs. High-dose ANS produces the same pattern, shifted towards the start codon and reduced in amplitude, indicating that both regimes follow a single dose-dependent positional rule rather than representing collision-positive and collision-negative states. At 30 min after UV irradiation, by contrast, collisions appear more dispersed and accumulate preferentially on highly translated mRNAs. These findings refine the interpretation of one of the most widely used experimental paradigms for inducing ribosome collisions and RSR signalling.

## Material and Methods

### Cell culture and treatments

Mouse embryonic stem cells (mESCs; male 129 × C57BL/6 background; same as in (15)) were cultured in DMEM supplemented with 15% fetal bovine serum (FBS), 1× non-essential amino acids, 1 mM sodium pyruvate, 2 mM L-glutamine, 0.1 mM β-mercaptoethanol, and 1× penicillin/streptomycin. Leukemia inhibitory factor (LIF) was added at a 1:1000 dilution from homemade stocks. Culture dishes were coated with gelatine prior to seeding. Cells were passaged every 2 days and maintained at 65-70% confluency. Before treatments, culture medium was refreshed 2 h before ANS/DMSO addition or UV irradiation. Cells were incubated with ANS (Sigma #A9789) or DMSO for 15 min at the indicated concentrations. For UV irradiation, medium was removed, the dish irradiated in a Vilber Lourmat BIO-SUN++ irradiation system (wave-length 312 nm; 0.05 J/cm^2^) and the original medium was immediately returned to the plate for the recovery phase.

### Western blot

To harvest cells for Western blot analysis, the dish was rinsed and cells were scraped down in cold PBS. After collecting cells by centrifugation, cells pellets were frozen in liquid nitrogen before protein extraction. Cells were lysed in RIPA buffer (150 mM NaCl, 50 mM Tris pH 8.0, 1% NP40, 0.5% sodium deoxycholate and 0.1% SDS) supplemented with complete protease and phosphatase inhibitors (Roche). Equal amounts of protein were resolved on bis-tris 4%- 12% gradient gels (Life Sciences) before transferring to PVDF membranes (iblot system, Life Sciences). Membranes were blocked for 1 h in TBST + 5% milk, followed by overnight incubation at 4 °C with primary antibodies against phospho-JNK (Cell Signaling Technology #4668S), Vinculin/VCL (Abcam #ab129002), RPS3 (ProteinTech #11990-1-AP), or phospho- eIF2alpha (Abcam #ab32157). After washing in TBST, membranes were incubated with secondary antibodies (Promega #W401B or #W402B) for 1 h at room temperature. Signal was developed using ECL (Thermo #34095) and detected using a Fusion-FX6 system.

### Polysome analysis

Polysome analyses were carried out on mESCs treated with the indicated conditions. ANS at 10 mg/L was included in all lysis and gradient buffers irrespective of treatment condition to minimise post-harvest changes in ribosome occupancy and collision status. Prior to lysis, cells were washed with ice-cold PBS and lysed directly on the plate in 250 µL of lysis buffer (50 mM Hepes-KOH pH 7.3, 100 mM potassium acetate, 15 mM Mg(OAc)₂, 5% glycerol, 0.5% Triton X-100, 2× complete protease inhibitor cocktail, 1× phosphatase inhibitor cocktail, 5 mM DTT, and 10 mg/L anisomycin). Lysates were cleared from debris by centrifugation at 9000 rpm for 10 min at 4 °C in a microfuge. Nuclease treatment prior to polysome analysis was conducted by supplementing the appropriate volume of lysate corresponding to 180 µg of protein (ca. 120-150 µL) lysate with 1 mM CaCl₂, 3 µL (6000 U) MNase (NEB M0247S) and incubation for 30 min at 22 °C. Reactions were stopped with 2 mM EGTA. Samples were then loaded on a 14 ml gradient (10%-40% sucrose in 5 mM Hepes pH 7.3, 20 mM KOAc, 1 mM Mg(OAc)_2_, 10 mg/L anisomycin); for non-digested samples, comparable amounts of clarified lysate (equivalent of 180 µg protein, similarly treated with CaCl_2_ and EGTA, but no MNase) were loaded as well. After ultracentrifugation of the gradients (Beckman SW40 rotor; 4 °C, 1 h 45 min, 40’000 rpm), fractionation of the gradients was carried out using a customised ÄKTA system (GE Healthcare). For every sample, biological replicates were produced (N=3 independent experiments)

### Sample collection for downstream mass-spec analysis

Relevant fractions from the sucrose gradients were collected, flash-frozen in liquid nitrogen and stored at -80 °C. After thawing, proteins were precipitated on ice with 10% TCA, collected by centrifugation in a microfuge (4 °C, 20’000 rpm, 30 min), and pellets were washed twice with ice-cold 100% acetone. Pellets were then briefly air-dried and processed at the in-house mass spectrometry facility, as described in the next section. For input samples (used for later normalisation), the same clarified protein lysates that were also used for the loading onto the sucrose gradients, were set aside.

### Mass-spec sample preparation and protein digestion

For input samples, proteins were prepared and digested following the SP3 protocol (16). Briefly, protein extracts were adjusted to 2% SDS, 10 mM DTT, 50 mM Tris-HCl (pH 7.5), then heated for 10 min at 75 °C. Iodoacetamide was added to a final concentration of 30 mM, and samples were alkylated for 45 min in the dark. Sera-Mag™ SpeedBeads (Cytiva, 45152105050250) were added at a 10:1 (w:w) bead-to-protein ratio, and proteins were precipitated on the beads using ethanol (final concentration 60%). After three washes with 80% ethanol, proteins were digested on-bead in 50 µL of 100 mM ammonium bicarbonate containing 0.6 µg trypsin (Promega, #V5113) for 1.5 h at 37 °C. The same quantity of trypsin was then added, and incubation continued for another 1.5 h. Supernatants were collected and mixed with two sample volumes of isopropanol containing 1% TFA, followed by desalting on an SCX plate (Oasis MCX 96-well, 186001830BA; Waters^TM^) to remove residual SDS. After washing with isopropanol/1% TFA and 2% acetonitrile/0.1% FA, peptides were eluted in 150 µL of 40% MeCN, 59% water, and 1% (v/v) ammonia, then dried by centrifugal evaporation. For polysomes samples, proteins were digested following a modified version of the iST method (17) named miST method. Samples were redissolved in 40 µl miST lysis buffer (1% Sodium deoxycholate, 30 mM Tris pH 8.6, 10 mM DTT), and heated for 10 min at 75 °C. They were then diluted 1:1 (v:v) with water containing 4 mM MgCl_2_ and benzonase (Merck #70746, 100× dilution of stock = 250 Units/µl), and incubated for 15 minutes at RT to digest nucleic acids. Reduced disulfides were alkylated by adding 0.25 volumes of 160 mM chloroacetamide (32 mM final) and incubation for 45 min at RT in the dark. Samples were adjusted to 3 mM EDTA and digested with 1.0 µg Trypsin/LysC mix (Promega #V5073) for 1 h at 37 °C, followed by a second 1 h digestion with an additional 0.5 µg of proteases. To remove sodium deoxycholate, two sample volumes of isopropanol containing 1% TFA were added to the digests, and the samples were desalted on a strong cation exchange (SCX) plate (Oasis MCX 96-well, 186001830BA; Waters^TM^) by centrifugation. After washing with isopropanol/1%TFA, peptides were eluted in 150 µl of 40% MeCN, 59% water, 1% (v/v) ammonia, and dried by centrifugal evaporation.

### Liquid Chromatography-Mass Spectrometry analyses

LC-MS/MS analyses were carried out on a TIMS-TOF Pro (Bruker, Bremen, Germany) mass spectrometer interfaced through a nanospray ion source (“captive spray”) to an EvoSep One liquid chromatography system (EvoSep, Odense, Denmark). Peptides were separated on a reversed-phase Aurora Elite C18 column (15 cm, 75 µm ID, 1.7 µm, IonOpticks) at a flow rate of 200 nL/min with a 20 sample per day method (runtime time: 68 min, solvents were water and acetonitrile with 0.1% formic acid).

Data-independent acquisition was carried out using a method similar to a standard DIA-PASEF method reported previously(18), with ion accumulation for 100 ms for each the survey MS1 scan and the MS2 scans. Duty cycle was kept at 100%. Precursor ions were chosen within a reduced mobility range from 1/k0 =0.7 to 1.4 and between m/z=400 to 1200. Collision energy was ramped linearly based uniquely on the 1/k0 values from 20 (at 1/k0=0.6) to 59 eV (at 1/k0=1.6). Per cycle, the mass range 400-1200 m/z was covered by a total of 20 windows, each 40 Th wide, with a total cycle time of 1.3 s.

### MS data processing

Identification of peptides directly from DIA data was performed with Spectronaut 20.4 with the Pulsar engine using the “deep” setting and searching the reference mouse proteome (www.uniprot.org) database of February 6^th^, 2025 (54’747 sequences), and a contaminant database containing the most usual environmental contaminants and enzymes used for digestion (from (19)). For identification, peptides of 7-52 amino acid (AA) length were considered, cleaved with trypsin/P specificity and a maximum of 2 missed cleavages. Carbamidomethylation of cysteine (fixed), methionine oxidation and N-terminal protein acetylation (variable) were the modifications applied. FDRs for peptide and protein group identifications were all at 1%. Ion mobility for peptides was predicted using a deep neural network and used in scoring. Peptide-centric analysis of DIA data was carried out using the library generated by Pulsar engine. Single hits proteins (defined as matched by one stripped sequence only) were kept in the Spectronaut analysis. Peptide quantitation was based on XIC area, for which a minimum of 1 and a maximum of 3 (the 3 best) precursors were considered for each peptide, from which the median value was selected. Quantities for protein groups were derived from inter-run peptide ratios based on MaxLFQ algorithm (20). A local regression normalisation was applied, as described by Callister et al. (21).

All subsequent analyses were carried out with an in-house developed software tool (available on https://github.com/UNIL-PAF/taram-backend). Contaminant proteins were removed, and quantity values for protein groups generated by Spectronaut were log_2_-transformed. After assignment to groups, only proteins quantified in all samples of at least one group were kept. Missing values were imputed based on a normal distribution with a width of 0.3 standard deviations (SD), down-shifted by 1.8 SD relative to the median.

### MS data normalisation and post-processing

Input and ribosome fraction protein mass spectrometry quantifications containing imputed data were filtered to retain a single protein per PG.Genes entry. Relative abundance was determined by normalising the quantity of each protein in a polysomal fraction to its quantity in the corresponding input sample. As additional normalisation, and to account for differences in total ribosome content across fractions, individual protein quantities were further expressed relative to median relative abundance (normalised to input, described above) of the ensemble of ribosomal proteins (using all detectable small and large subunit proteins).

For all statistical analyses log_2_-transformed data was used. Principal component analysis (PCA) was performed on the normalised abundances to evaluate overall sample clustering. Differential protein abundance between control and treated conditions was evaluated using two-sample two-sided *t*-tests without FDR correction and with a nominal p-value cut-off of 0.05. These values are reported in **Supplementary Table S1**.

### Ribosome profiling

Ribo-seq was carried out on mESCs treated with LD ANS (0.1 mg/L anisomycin), HD ANS (100 mg/L anisomycin) or UV (30 min recovery post-UV), or control-treated (i.e. DMSO for ANS condition; UV-free for UV condition). The same collection and lysis conditions were applied to all treatment and control samples. Cells were rapidly rinsed and immediately collected in ice-cold PBS supplemented with 200 µg/mL cycloheximide (CHX). Cell lysis was performed according to our standard protocol (22) and with the same lysis buffer, i.e. 150 mM NaCl, 20 mM Tris-HCl pH 7.4, 5 mM MgCl_2_, 5 mM DTT, 1% Triton X-100, 0.5% Sodium deoxycholate, 100 µg/mL cycloheximide, complete EDTA-free protease inhibitors (Roche), 40 U/mL RNasin plus (Promega), then cleared from debris, quantified via OD260. Ribosome-protected fragments (RPFs) were generated by digesting lysates with RNase I (Ambion; 43 U/OD) in the presence of Turbo DNase (Ambion; 0.3 U/OD). Reactions were inactivated with SUPERase·In™ RNase Inhibitor (8.7 µL; Invitrogen) and RPF-containing ribosomes were enriched on S-400 HR columns (Cytiva) before RNA extraction from the flow-through using the miRNeasy kit (Qiagen). 2.5 µg of purified RNA was separated on 15% TBE-urea gels, and RPFs corresponding to the monosome and disome footprint lengths were excised from the gels and purified as described. Library preparation followed essentially the protocol by McGlincy & Ingolia (23), with minor modifications. Briefly, RNA fragments were 3′-repaired using PNK, followed by ligation of 5′-adenylated adapters containing sample barcodes and an 8-nt UMI using T4 RNA Ligase 2 (truncated KQ). Samples were multiplexed, and adaptor removal was performed for 1 h at 30°C and 1 h at 37°C by treating libraries with 2 µl of a 1:1 mix of 5′ deadenylase (NEB) and RecJf (NEB). rRNA depletion was carried out using a customised siTools rRNA depletion kit, with subsequent purification using Zymo RNA Clean & Concentrator columns. cDNA synthesis was performed with EpiScript (Biosearch Technologies), followed by sequential clean-up steps using Exonuclease I and a 1:1 mix of RNase I and Hybridase. Circularized cDNA libraries were generated using CircLigase II, and PCR amplification (Phusion polymerase, NEB) was performed using i5 (Nextera D501, D502, D503, D504, D505 or D507) and i7 (#iA_01, #iA_02, #iA_03, #iA_04, #iA_05 or #iA_08) indexing primers. Final libraries were sequenced in single-end mode on an Aviti sequencer.

For overall batch structure of cell harvesting, library generation and sequencing, LD ANS libraries (treatment and controls) were produced in one batch (3 control, 3 treated biological replicates for monosomes and disomes; as one treated monosome replicate and one control disome replicate showed very low read numbers in the final library, these were omitted from the analyses), and the remaining samples (HD ANS, UV, and corresponding controls) together as a second batch (4 biological replicates for treatments and 4 for controls). Co-clustering of Ribo-seq control samples across datasets in the PCA validates reproducibility across batches.

### RNA-seq datasets and analysis

Complementary RNA-seq datasets were produced from RNA extracted from aliquots of the same lysates that were used for footprint generation (n=3 biological replicates each for LD ANS and its control; n=4 biological replicates each for HD ANS and its control as well as UV and its control). Total RNA was extracted with Qiazol and libraries were generated following Illumina Stranded Total RNA Prep with Ribo-Zero Plus protocol and were single read sequenced on Aviti sequencer. Raw sequencing reads were trimmed using TrimGalore (v. 0.6.10; adapter="CTGTCTCTTATACACATCT" and minimum length=50 nt) and mapped to the mouse genome (*Mus musculus* GRCm39.111) using STAR (v.2.7.11b; parameters: -- seedSearchStartLmax 50 --outSAMmultNmax 1 --outSAMtype BAM SortedByCoordinate Unsorted --outReadsUnmapped Fastx --alignSJDBoverhangMin 1 --alignIntronMax 1000000 --outFilterType BySJout --alignSJoverhangMin 8 --sjdbGTFfile Mmusculus.GRCm39.111.sorted.gtf). We next generated read counts (in CDS) per transcript using the function summarizeOverlaps() (mode=Union; package GenomicAlignments in R; v4.4.0) and performed differential gene analysis using DESeq2 (v. 1.46.0).

### Monosome and Disome footprint sequencing data processing

To make our previous Ribo-seq pipeline (6) more user friendly, we integrated the commands in an automated and portable Snakemake workflow and included new tools. First, we removed adapters from the sequencing reads using TrimGalore (v0.6.10; auto-detect adapter type and minimum length = 30 nt). We filtered the reads according to length (monosomes 26-37 nt; disomes 43-68 nt) and applied a quality filter using fastq_quality_filter (fastx_toolkit v0.0.14). The remaining reads were sequentially mapped using STAR (24) (v2.7.11b; parameters: : -- seedSearchStartLmax 28 --outSAMmultNmax 1 --outSAMtype BAM SortedByCoordinate Unsorted --outReadsUnmapped Fastx) on mouse (*Mus musculus* GRCm39.111) rRNA, human (*Homo sapiens* GRCh38.111) rRNA, and mouse tRNA. The unmapped reads were demultiplexed (UMI-tools whitelist/extract; v1.0.0git (25); and demuxbyname.sh from BBTools at https://bbmap.org/ to split reads by barcode) and mapped to the mouse genome (parameters: --outSAMunmapped Within --outFilterType BySJout --outSAMattributes NH HI AS NM MD --outFilterMultimapNmax 20 --outFilterMismatchNmax 999 --alignIntronMin 20 -- alignIntronMax 1000000 --alignMatesGapMax 1000000 --alignSJoverhangMin 8 -- alignSJDBoverhangMin 1 --sjdbScore 1 --genomeLoad NoSharedMemory --outSAMtype BAM Unsorted --quantMode TranscriptomeSAM). Genome-aligned reads were then projected to the transcriptome and filtered to keep only deduplicated reads (UMI-tools dedup) mapped to the forward strand. Next, we obtained read counts (in CDS) per transcript using the function summarizeOverlaps() (mode=’IntersectionNotEmpty’; package GenomicAlignments in R; v4.4.0) (26) and selected reads mapped to the single most expressed transcript per protein-coding gene. The reads mapped multiple times within a transcript were kept only once and a single histone representative per family (19 transcripts) was selected to reduce the number of multimapped reads in the histone families. Moreover, reads mapping to *Lars2*, *Grcc10*, and *Ftl1* were excluded prior to downstream analysis, because we noted that these loci produced abundant reads from UTR regions that according to our visual analysis did not correspond to ribosome footprints but to RNA hairpin structures that are resistant RNaseI treatment. To obtain the P-site offset per read length for monosome data, we used the function psite() (parameters: flanking=6, extremity=’5end’; package riboWaltz in R (27)) and added 3 nt. Due to the lack of optimised A-site offset estimation methodology for disome data, we defined the A-site offset of the leading ribosome as the read 5’ end position +45 nt (i.e., a simplified approach to that of (6)). For each transcript, we then obtained the number of A-site counts per position and converted these values into codon counts.

To identify ribosome pausing sites we adapted the methodology from (28), comparing the codon counts at each position of a transcript with the codon counts in the CDS (minus the count at the investigated position) between treated and non-treated samples. To do so, we created 2x2 contingency tables and performed two-tailed Fisher’s exact tests to identify changes in occupancy at each codon position. Sites were labelled as “up”, “down”, or “ns” based on their fold-change and statistical significance. To determine whether specific codons were significantly enriched or depleted among target sites compared to expected background distributions, the 1000 most significant up-regulated sites (“up”) were compared to non-significant (“ns”) sites within the same gene set by random permutation (n = 10’000). Site labels (“up” vs. “ns”) were randomly shuffled to compute null distributions of codon counts among target sites. Observed codon frequencies (obs) were evaluated against permutation-derived expectations (exp_mean and exp_sd) to calculate standard Z-scores (Z=(obs- exp_mean)/exp_sd). Two-sided empirical p-values were then computed to identify significantly enriched or depleted codons (p<=0.05).

### Nucleotide resolved data processing and analysis

Using the nucleotide-level counts table (.sga), raw data were either variance-stabilizing transformed (VST) for Principal Component Analysis (PCA) or expressed as counts per million for metagene profiling. For the metagene analysis, footprint positions were calculated relative to annotated start and stop codons. The mean normalised counts per condition were then summed at each nucleotide position to generate metagene plots.

### Fold-change metagene plots

For the visualisation and analysis of the FC-based metagene plots, SGA files were converted into BigWig files using the sga_to_bigwig.py script. Subsequent processing was primarily carried out using the deepTools suite (29). Replicates were averaged for each condition using the bigwig_mean.py script (--binSize 1). Fold-change density files were generated using the bigwig_compare.py script (--operation log2 --binSize 5 --pseudocount 0.1 0.1). Metagene profiles and heatmaps were generated using computeMatrix.py, followed by plotProfile.py or plotHeatmap.py, respectively. Metagene profiles were subsequently formatted and analyzed using GraphPad Prism. To select for a robust set of disome-containing transcripts in these comparisons, we restricted analyses to the 5000 transcripts with the highest disome signal across the CDS under control treatment (based on the DMSO dataset of the LD ANS treatment experiment). These transcripts were grouped by k-means clustering based on their fold-change profiles in the UV condition across the −100 to +200 nt region relative to the translation start site. The list of the 5000 selected transcripts is provided in **Supplementary Table S2**. Gene Ontology analyses were performed using the ShinyGO web application (30).

### Codon dwell times (Ribo-DT)

Codon dwell times at the E-, P- and A-sites were estimated with Ribo-DT (31). Deduplicated reads aligned to the transcriptome were realigned to the mouse genome (GRCm39, Ensembl release 111) with STAR 2.7.0f, and only uniquely mapped reads without mismatches were used. A-site assignment: For monosomes, footprints of 26-35 nt were used, and the A-site offset was determined separately for each read length and frame from the distribution of 5′ ends around annotated start codons, assuming the start codon in the P site. For disomes, footprints of 51-68 nt were used, and the A site of the leading ribosome was placed 15 nt upstream of the read 3′ end for all lengths. Model: Only single-codon terms were fitted (no codon-pair interaction terms). To include the post-initiation zone, which the original implementation excludes (it requires a full 40-codon window around the A site, i.e. 75 nt upstream), each complete coding sequence was extended with 75 nt of 5′ UTR and 45 nt of 3′ UTR, so that A sites from the second codon to the stop codon were modelled. Window positions outside the coding sequence were not assigned a codon identity but were modelled by position-specific indicator variables. Statistics: Monosome libraries were fitted per biological replicate, and treated and control dwell times were compared for each codon and site with a two-sample t-test, with Benjamini-Hochberg (BH) correction within each condition and site. Codons with an adjusted p-value below 0.05 and an absolute change above 0.5 log2 were considered significant, and those whose change is also an outlier among the 61 codons (|robust z| > 2.5) are labelled. Several individual disome libraries were too shallow for the model to converge, so disome replicates were pooled per condition, and changes were assessed with a z-test on the model standard errors. Because pooled libraries carry counting noise only, codons were considered significant only if their change was also an outlier among the 61 codons (|robust z| > 3).

## Results

### Establishing anisomycin and UV conditions that induce ribosome collisions in mESCs

To compare drug- and UV-induced ribosome impairment, we established low-dose (LD) ANS, high-dose (HD) ANS and UVB conditions in mESCs, each with matched controls, and analysed them by polysome profiling, Ribo-seq, RNA-seq and proteomics (schematic in **Fig. 1A**).

**Figure 1.**
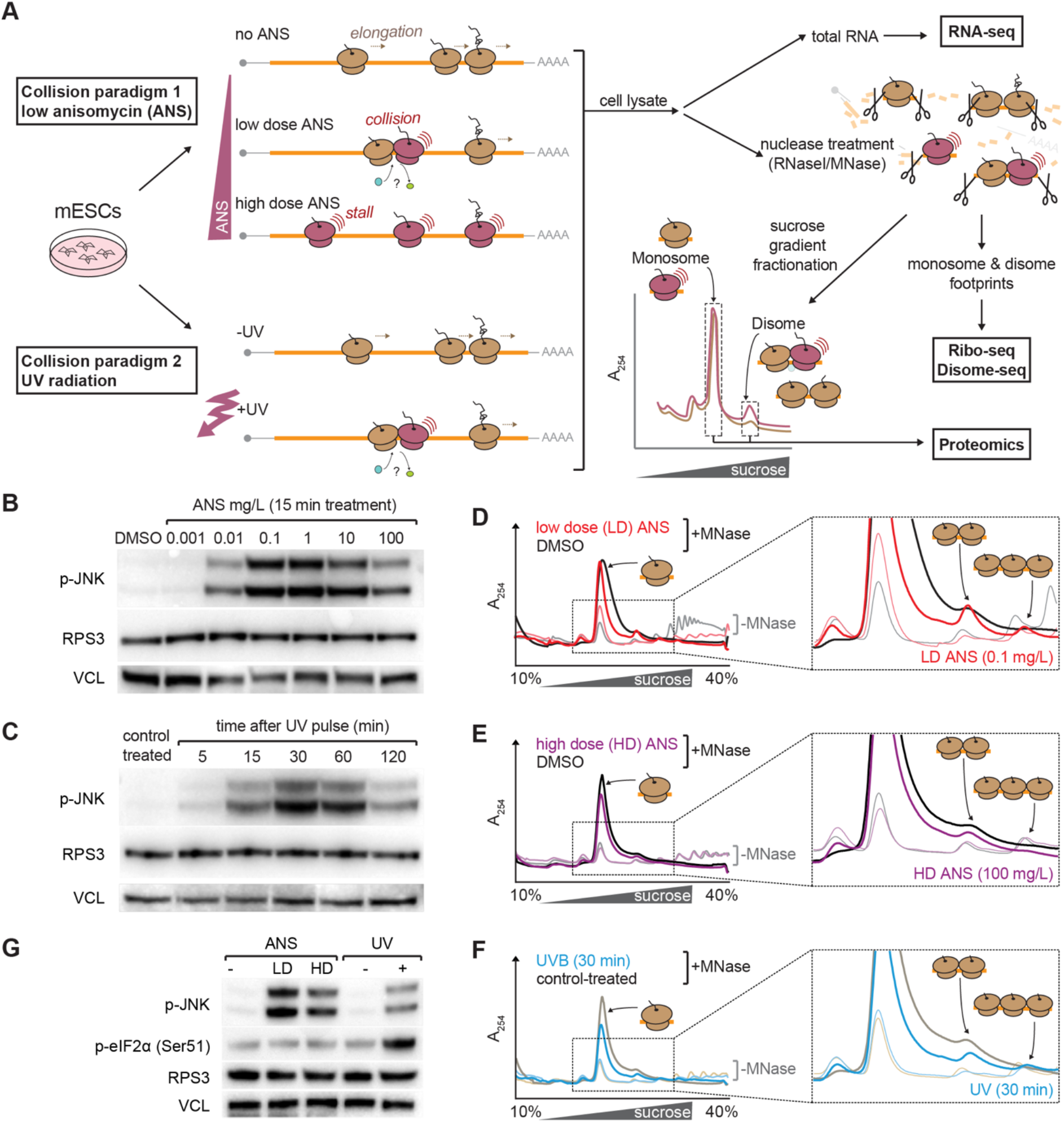
Defining anisomycin and UV regimes that trigger ribosome collisions and RSR activation in mESCs. **A.** Schematic overview of models for the occurrence of ribosome collision and of the experimental workflow used to characterize collision events in this study. mESC lysates were subjected to sucrose gradient fractionation with or without nuclease treatment (RNase I or MNase), followed by RNA-seq, Ribo-seq, disome profiling, and proteomic analyses. **B.** Western blot analysis of JNK phosphorylation levels in mESCs treated with increasing concentrations of anisomycin (ANS) for 15 min. VCL (Vinculin) and RPS3 are shown as loading controls. **C.** Western blot analysis of JNK phosphorylation levels in mESCs harvested at the indicated time points following UV irradiation (0.05 J/cm²). VCL and RPS3 serve as loading controls. **D.** Representative polysome profiles of mESCs treated with low-dose (LD) ANS (0.1 mg/L, 15 min) or DMSO as control. Polysome-containing extracts were digested with MNase (dark traces) before loading onto gradient, to collapse polysomes into monosomes, disomes, and higher-order ribosome collisions. Non-digested extracts are show as light traces. **E.** As in panel D, following high-dose (HD) ANS treatment (100 mg/L, 15 min). **F.** As in panel D, UV-treated mESCs (0.05 J/cm², 30 min post-irradiation) and corresponding control-treated cells. **G.** Comparative analysis of JNK and eIF2α phosphorylation under LD ANS, HD ANS and UV treatment. Independent samples from panels B and C were used. Alt text: Seven panels. A is a schematic: a dish of mESCs branches into two collision paradigms, low-dose anisomycin and UV radiation, each drawn as cartoon mRNAs carrying tan elongating ribosomes, with dark red arrested ribosomes and red radiating lines marking collisions, and a wedge indicating increasing anisomycin dose; to the right, cell lysate branches to total RNA for RNA-seq, to nuclease treatment with RNase I or MNase yielding monosome and disome footprints for Ribo-seq and disome-seq, and to sucrose gradient fractionation, drawn as an absorbance trace with labelled monosome and disome peaks, feeding into proteomics. B and C are western blot panels with rows for phosphorylated JNK, RPS3 and vinculin: B across a DMSO control and six anisomycin concentrations from 0.001 to 100 mg/L, C across a control and five times from 5 to 120 minutes after UV. D, E and F each show sucrose gradient absorbance traces plotted against a 10 to 40 % sucrose gradient, with nuclease-digested traces bold and undigested traces pale, treated in colour over a black or grey control, and a boxed inset magnifying the monosome, disome and trisome region with ribosome cartoons marking each peak; D is low-dose anisomycin, E high-dose anisomycin, F UVB at 30 minutes. G is a western blot with lanes for untreated, low-dose and high-dose anisomycin and for untreated and UV-treated cells, with rows for phosphorylated JNK, phosphorylated eIF2-alpha at serine 51, RPS3 and vinculin.

We first titrated ANS (15 min) and monitored JNK phosphorylation, a commonly used readout of RSR activation (10,32) (**Fig. 1B**). Phospho-JNK became detectable at 0.01 mg/L and was maximal at 0.1-1 mg/L. We defined 0.1 mg/L as the LD condition, in line with studies in other cell types (9). For the HD condition we chose 100 mg/L anisomycin, a concentration commonly used to arrest translation rapidly (33,34). At this dose, phospho-JNK was reduced relative to LD ANS but remained clearly detectable (**Fig. 1B**). This differs from reports that high ANS concentrations abolish JNK activation in other systems (10), and indicated that strong elongation inhibition in mESCs attenuates rather than eliminates this response.

We then assessed collided ribosomes by sucrose-gradient analysis of MNase-digested lysates, in which nuclease-resistant disomes and trisomes report on collisions (schematic in **Fig. 1A**, right). To limit post-lysis changes in ribosome states, all samples were lysed and fractionated in the presence of 10 mg/L ANS (see Materials and Methods). LD ANS markedly increased disome and trisome peaks relative to DMSO-treated controls (**Fig. 1D**). HD ANS also produced disomes, though at lower levels than LD ANS (**Fig. 1E**). Thus HD ANS reduced, but did not eliminate, collided ribosomes, consistent with the JNK data.

For UV, we exposed cells to a UVB pulse (0.05 J/cm^2^) as previously described (12) and followed recovery from 5 to 120 min. JNK phosphorylation increased progressively, peaked at 30 min and declined thereafter (**Fig. 1C**). At 30 min, nuclease-resistant disomes and trisomes were detectable (**Fig. 1F**), and we used this time point for all subsequent experiments.

Finally, we compared JNK and eIF2α phosphorylation, as proxies for RSR- and ISR-associated signalling, respectively, across conditions (**Fig. 1G**). LD ANS induced stronger JNK phosphorylation than UV and did not increase eIF2α phosphorylation. UV, in contrast, induced robust eIF2α phosphorylation, consistent with GCN2-mediated ISR activation after UV (14). ISR activation is expected to reduce translation initiation, a point that becomes relevant for interpreting the UV ribosome profiling data below.

Together, these experiments define three conditions that induce JNK phosphorylation and collided ribosome species to different extents, and that differ in ISR engagement. They provide the basis for comparing where collisions form and which factors associate with them.

### Collided-ribosome fractions show shared and condition-specific co-sedimentation of surveillance and initiation factors

To examine which factors associate with monosomes and collided ribosomes under each condition, we analysed monosome and disome fractions from sucrose gradients of MNase-digested lysates by quantitative mass spectrometry (schematic in **Fig. 1A**, right). Protein abundances in each fraction were normalised to the corresponding input and then to the median abundance of ribosomal proteins (RPs) in that fraction. This approximates protein abundance per ribosome and reduces effects of differences in the amount of ribosomal material recovered (**Supplementary Fig. S1A; Supplementary Table S1)**. Treatment- associated changes were most pronounced after UV, where treated and control samples separated clearly in principal component analysis; separation was weaker for the ANS comparisons (**Fig. 2A-C; Supplementary Fig. S1B**). Because of this, and because statistical testing used nominal p-values (n = 3), we focused on predefined sets of factors with known roles in ribosome surveillance and translation, and placed weight on changes that were consistent across fractions or conditions or shared by several components of the same complex.

**Figure 2.**
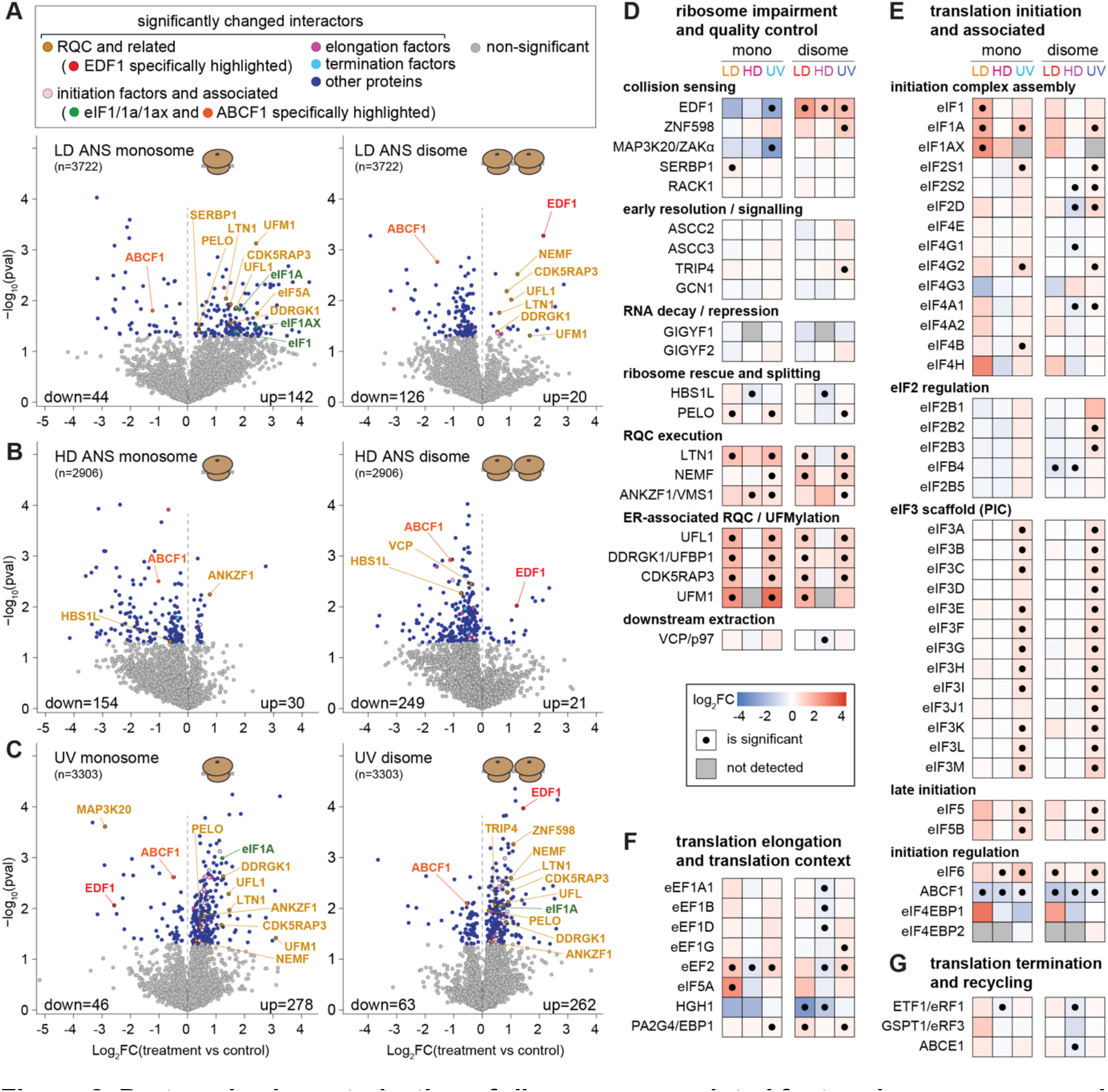
Proteomic characterization of ribosome-associated factors in monosome and disome fractions under ANS and UV conditions. **A.** Volcano plots showing protein abundance changes in monosome fractions (left panel) and disome fractions (right panel) following low-dose (LD) ANS treatment compared to control. Highlighted proteins include factors involved in ribosome-associated quality control (RQC), initiation, and elongation. Number of detected proteins and counts of up- and downregulated proteins are indicated. Data normalisation per sample takes into account input abundance of the proteins, as well as a normalisation step relative to the median abundance of detected ribosomal proteins (RPs) within each fraction, allowing interpretation of changes on a per-ribosome basis. N=3 per condition. Nominal p-values are from *t*-test and are also reported in **Supplementary Table S1**. **B.** As in panel A, for monosome and disome fractions following HD ANS treatment. N=3 per condition. **C.** As in panel A for monosome and disome fractions following UV treatment compared to control. N=3 per condition. **D.** Functional categorisation of proteins associated with ribosome impairment and quality control pathways, including collision sensing, ribosome rescue and splitting, RQC execution, ER-associated RQC/UFMylation, and downstream extraction. Proteins are grouped by function and annotated according to their roles in translation and ribosome surveillance. The heatmap indicates the log_2_FC, and a dot in a box statistical significance (nominal p < 0.05; t-test; see **Supplementary Table S1**). **E.** Enrichment of translation initiation factors and associated complexes across monosome and disome fractions under LD ANS, HD ANS, and UV conditions. Selected components of different initiation complexes are indicated. Colouring and statistical significance as in panel D. **F.** As in panel E, for translation elongation-associated factors in monosome and disome fractions across conditions. Selected elongation factors and translation context regulators are shown. Colouring and statistical significance as in panel D. **G.** Enrichment of translation termination and ribosome recycling factors in monosome and disome fractions across conditions. Selected termination and recycling factors are indicated. Colouring and statistical significance as in panel D. Alt text: Seven panels. A, B and C are pairs of volcano plots, monosome on the left and disome on the right, for low-dose anisomycin, high-dose anisomycin and UV respectively. Each plots log_2_fold-change of treatment versus control on the horizontal axis against negative log_10_p- value on the vertical axis; most points are grey, significantly changed proteins are blue, and selected factors are labelled in colour according to a key above panel A that distinguishes RQC and related factors, initiation factors and associated proteins, elongation factors, termination factors and other proteins. Counts of down- and up-changed proteins are printed in the lower corners of each plot. D to G are heatmap grids in which rows are individual proteins grouped under functional sub-headings and the six columns are monosome and disome fractions for each of the three treatments; cells are shaded on a blue-to-red log_2_fold-change scale, a dot marks statistical significance and grey marks proteins not detected. D covers ribosome impairment and quality control, from collision sensing through RQC execution to downstream extraction; E covers translation initiation and associated complexes, grouped into initiation complex assembly, eIF2 regulation, the eIF3 scaffold, late initiation and initiation regulation; F covers translation elongation; G covers translation termination and recycling.

Among collision-associated factors, the sensor EDF1 was significantly enriched in disome fractions under all three conditions, to a similar extent, consistent with its role as a general collision sensor (35) (**Fig. 2D**). Under UV, EDF1 was concomitantly depleted from monosome fractions, suggesting redistribution towards collided species. The E3 ligase ZNF598 (9) was significantly enriched only in UV disomes. The RSR kinase ZAKα (MAP3K20) was depleted from monosome fractions, most strongly under UV, but largely unchanged in disome fractions, whereas its ribosomal binding partner RACK1 was unchanged throughout. SERBP1, which has been reported to compete with ZAKα for binding to RACK1 (7), showed a modest trend towards enrichment that reached nominal significance only in LD ANS monosomes, and no consistent pattern emerged across conditions.

Several factors acting downstream of collision sensing were also enriched, including the rescue factor PELO (36) (modestly) and the RQC components LTN1 (37), NEMF (38) and ANKZF1 (39), most consistently under UV (**Fig. 2D**). Components of ER-associated RQC and the UFMylation machinery, UFL1, DDRGK1, CDK5RAP3 and UFM1 (40), were enriched as a group under both LD ANS and UV, suggesting that a subset of impaired ribosomes is engaged in membrane-associated translation. In contrast, VCP/p97 (41), components of the ASC-1 complex (42,43), GCN1 (44) and GIGYF1/2 (45) showed little or no change. The captured complexes thus include ribosomes engaged by collision sensors and RQC factors, whereas factors acting at the extraction step, or in alternative pathways, were not detectably enriched.

Beyond quality-control factors, the most coherent change was an enrichment of translation initiation factors, which was most prominent under UV and included most eIF3 subunits as well as eIF2, eIF4G and eIF5 components in both monosome and disome fractions (**Fig. 2E**). Elongation (**Fig. 2F**) and termination factors (**Fig. 2G**) showed fewer and less consistent changes. Initiation factors can remain associated with 80S ribosomes during early elongation (46) and their enrichment is therefore consistent with impairment occurring shortly after initiation. Under UV, however, ISR activation (**Fig. 1G**) may also favour accumulation of 40S- containing initiation intermediates that co-sediment with these fractions – the substrate of initiation RQC (iRQC), in which RNF10 ubiquitylates uS3 and uS5 on 40S subunits whose progression past the start codon is blocked, including after ISR activation (47). The two possibilities cannot be distinguished from these data alone.

Finally, the initiation-associated ATPase ABCF1 (48,49) was depleted from both monosome and disome fractions under all three conditions (**Fig. 2E**), consistent with impaired ribosomes losing factors associated with productive translation.

Together, these data show that collided ribosomes under all three conditions recruit EDF1, that engagement of ZNF598 and downstream RQC factors is most evident after UV, and that initiation factors co-sediment with impaired ribosomes, an indication that ribosome impairment may be concentrated near the start of coding sequences.

### Low- and high-dose anisomycin produce a post-initiation collision zone whose position depends on dose

We next sought to map where ribosomal stalls and collisions occur across the transcriptome under the three impairment conditions. To this end, we generated monosome- and disome- resolved ribosome profiling libraries, together with RNA-seq libraries, from LD ANS-, HD ANS- and UV-treated cells and matched controls (**Supplementary Fig. S2A-J**). We first aligned footprints to start and stop codons using their inferred A-sites (for disomes, the A-site of the leading ribosome; see Materials and Methods) and compared metagene profiles between treated and control samples.

Under LD ANS, both monosome and disome footprints accumulated strongly near the start of coding sequences (**Fig. 3A, C**), with monosome signal peaking ∼30 nt and disome signal ∼63 nt downstream of the start codon (**Fig. 3E**). Cumulative distributions showed that the treatment-induced gain in footprint signal was largely confined to this region (**Fig. 3B, D**), whereas disome signal further downstream did not increase relative to control (**Fig. 3C**). Thus, under LD ANS, the additional collided ribosomes are not distributed along coding sequences, as commonly depicted (e.g. **Fig. 1A**), but are concentrated in a post-initiation zone.

**Figure 3.**
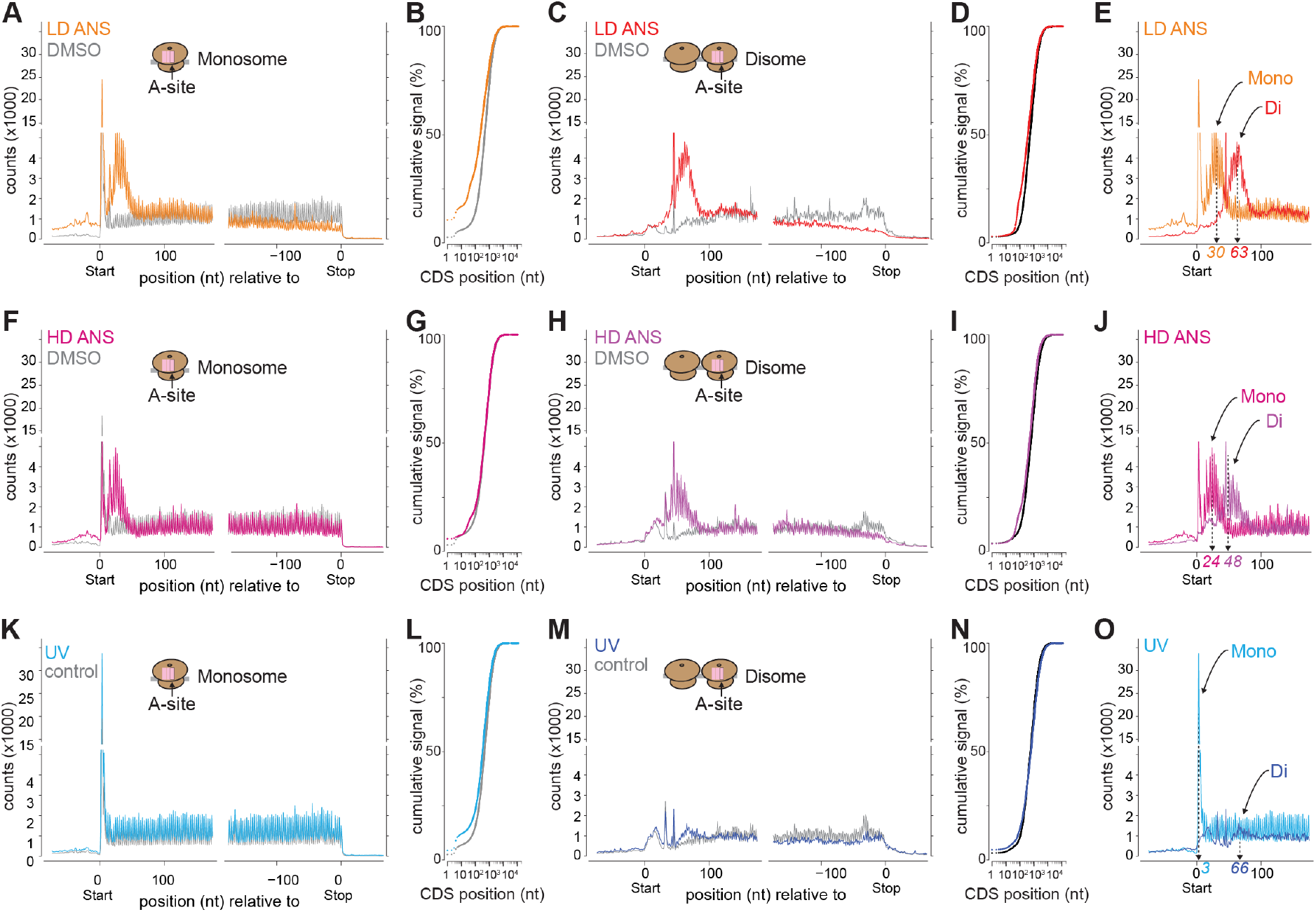
Metagene analysis of ribosome occupancy and collision positioning under ANS and UV conditions. **A.** Metagene profiles of monosome footprints, aligning inferred A-sites to start and stop codons, for mESCs treated with low-dose (LD) ANS (0.1 mg/L, 15 min) or DMSO control. Read densities are plotted relative to coding sequence (CDS) features start and stop. **B.** Cumulative monosome footprint signal along CDS positions under LD ANS and control conditions. **C.** Metagene profiles as in panel A, for disome footprints under LD ANS and control conditions. **D.** As in panel B, showing cumulative disome signal along CDS positions under LD ANS and control conditions. **E.** Overlay of monosome and disome LD ANS metagene profiles from panels A and C, allowing to visualise relative positions of peaks. **F.** Metagene profiles of monosome footprints as in panel A, but under high-dose (HD) ANS (100 mg/L, 15 min) and control conditions. **G.** Cumulative monosome footprint signal along CDS positions as in panel B, but under HD ANS and control conditions. **H.** Disome metagene profiles as in panel C, under HD ANS and control conditions. **I.** As in panel G, showing cumulative disome signal along CDS positions under HD ANS and control conditions. **J.** Analogous to panel E, overlay of monosome and disome HD ANS metagene profiles from panels F and G, allowing to visualise relative positions of peaks. **K.** Metagene profiles of monosome footprints following UV irradiation (0.05 J/cm², 30 min post- irradiation) and control treatment. **L.** Cumulative distribution of monosome footprint signal along CDS positions as in panel B, but under UV and control conditions. **M.** Metagene profiles of disome footprints following UV treatment and control conditions. **N.** As in panel L, showing cumulative distribution of disome signal along CDS positions under UV and control conditions. **O.** Overlay of monosome and disome UV metagene profiles from panels K and M. Alt text: Fifteen panels arranged in three rows, one per treatment: low-dose anisomycin, high- dose anisomycin and UV. Each row contains five plots. The first and third are metagene traces of A-site footprint counts against nucleotide position, split into a start-codon-aligned and a stop-codon-aligned window, with the treated sample in colour overlaid on a grey control; the first shows monosomes, the third disomes. The second and fourth are cumulative signal curves, rising from 0-100 % against CDS position on a logarithmic axis, with treated and control curves overlaid. The fifth superimposes the monosome and disome traces over the first hundred nucleotides, with dashed arrows marking the two peak positions numerically: 30 and 63 nucleotides for low-dose anisomycin, 24 and 48 for high-dose anisomycin, and 3 and 66 for UV.

HD ANS is often regarded as a collision-poor condition. Yet it produced the same pattern: monosome and disome peaks at ∼24 and ∼48 nt, closer to the start codon than under LD ANS (**Fig. 3F, H, J**), with an early-CDS gain in disome signal of lower amplitude (**Fig. 3G, I**). This agrees with the reduced but readily detectable JNK phosphorylation (**Fig. 1B**) and reduced levels of MNase-resistant disomes (**Fig. 1E**) under HD ANS. LD and HD ANS therefore do not represent collision-positive and collision-negative states. Rather, they produce the same positional pattern, shifted and attenuated as the dose increases.

The UV profile was different. Monosome footprints showed an increased peak at the start codon but little additional enrichment along the CDS (**Fig. 3K, L**), and disome footprints showed only a modest, broad increase at ∼66 nt (**Fig. 3M-O**).

A simple model can account for the ANS patterns. During the 15-min treatment, initiation continues while each elongation cycle carries a dose-dependent probability of drug-induced arrest. Ribosomes arrested within roughly one ribosome footprint (∼30 nt) of the start codon continue to occlude the initiation site and block further initiation; they appear as the monosome peak. Ribosomes that advance further before arresting vacate this region, allowing a new ribosome to initiate and run into them, which creates a disome. At higher doses, arrest occurs earlier, so both peaks shift towards the start codon and fewer ribosomes advance far enough to permit a second initiation, reducing the amplitude of the collision signal. Under UV, by contrast, ribosome impairment is accompanied by ISR activation (**Fig. 1G**), which is expected to reduce translation initiation and thus the supply of trailing ribosomes. This offers one plausible explanation for the comparatively weak and diffuse disome signal under UV.

### ANS produces a uniform collision zone across transcripts, whereas the UV response is heterogeneous

Metagene profiles are dominated by abundant, highly translated mRNAs. We therefore asked whether the ANS-induced collision zone is shared across transcripts or driven by a subset, and whether the modest UV effects reflect weak but widespread changes or stronger changes on fewer transcripts. To this end, we calculated position-resolved fold-changes (FC, treatment vs. control) for individual transcripts, restricting the analysis to the 5,000 transcripts with the highest disome coverage in control cells (**Supplementary Table S2**), and displayed them as aggregate profiles (**Fig. 4A, B**) and heat maps (**Fig. 4C, D**).

**Figure 4.**
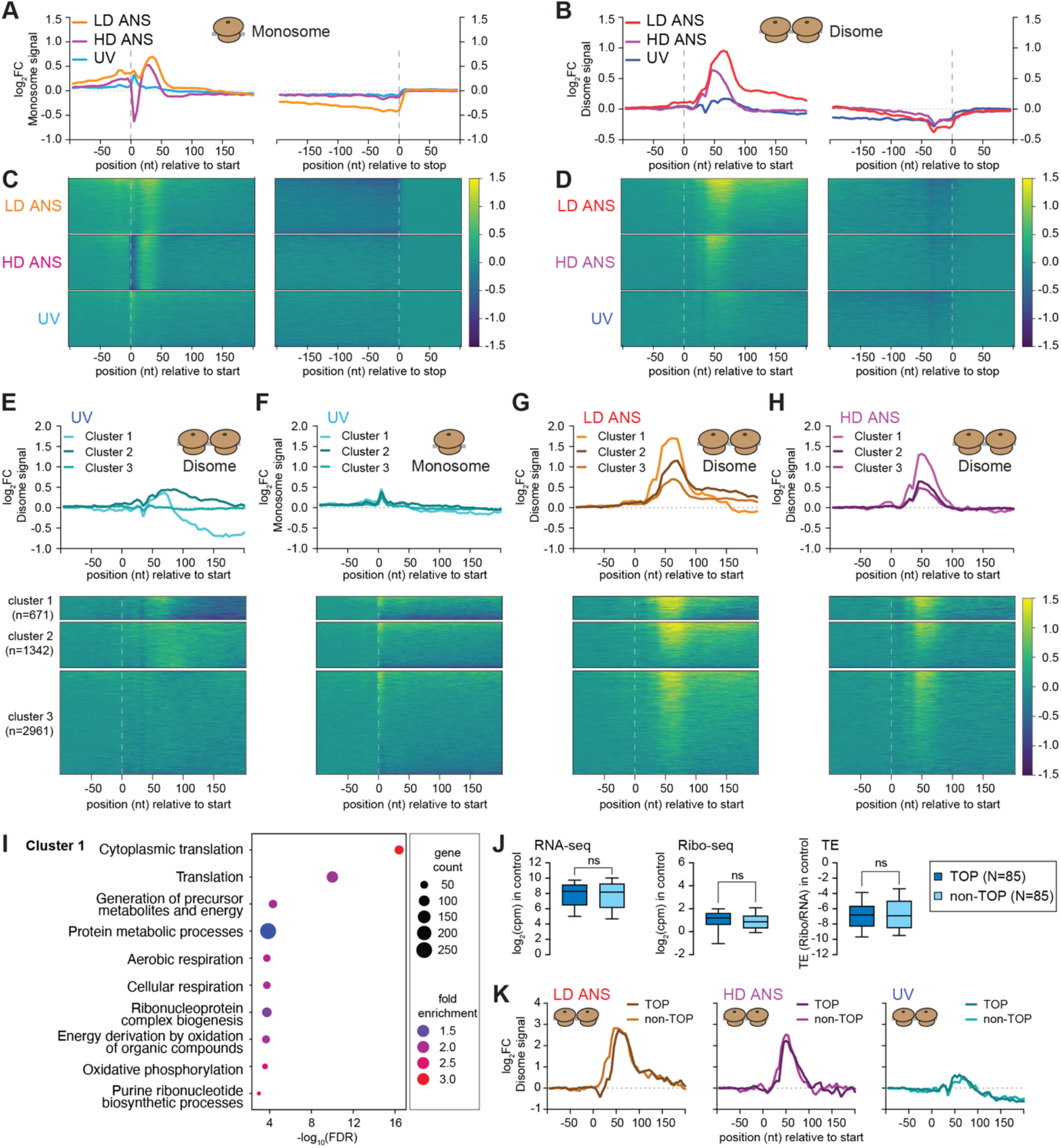
Anisomycin and UV induce distinct transcript-level collision landscapes. A. Aggregate profiles of position-resolved monosome footprint fold changes (log_2_FC, treatment versus control) across coding sequences. Individual transcript profiles were calculated using 5 nt-window occupancy measurements and subsequently aligned relative to start and stop codons. Profiles are shown for low-dose (LD) ANS, high-dose (HD) ANS, and UV conditions. The top-5000 highest disome transcripts were used in the analysis. B. As in panel A, showing disome footprint fold changes. C. Heat-map representation of monosome footprint fold changes across individual transcripts. Transcripts were ordered according to the magnitude of overall log_2_FC signal and aligned relative to start codons. Each row represents one transcript (N=5000 transcripts, as in panel A). D. As in panel C, showing disome footprint fold changes. Note the widespread post-initiation enrichment observed under LD and HD ANS conditions and the increased heterogeneity of UV-induced responses. E. K-means clustering of UV-induced disome fold-change profiles. Upper panels show metagene representations of the three clusters. Lower panels show the corresponding transcript heat maps. Numbers of transcripts per cluster are indicated. Transcript order in clusters follows decreasing fold-change signal. F. UV-induced monosome fold-change profiles for the same clusters as in panel E, with metagene profiles (top) and corresponding heat maps (bottom). Transcript order in clusters follows decreasing fold-change signal (i.e., different order as in panel E). G. Disome fold-change profiles of transcripts assigned to the UV-derived clusters shown in panel E, analysed under LD ANS conditions. Cluster-average metagene profiles (top) and heatmaps (bottom) are displayed. Transcript order in clusters follows decreasing fold-change signal. H. As in panel G, showing the same UV-derived transcript clusters analysed under HD ANS conditions. Transcript order in clusters follows decreasing fold-change signal. I. Gene ontology enrichment analysis of transcripts belonging to the collision-enriched UV cluster 1. Top-scoring significant biological process categories are shown. J. Boxplots depicting steady-state RNA-seq, Ribo-seq, and TE levels in control samples (from LD ANS series) for 5′ TOP mRNAs and a matched set of non-5′ TOP mRNAs. The non-5′ TOP matched set was generated using a nearest-neighbour approach. The absence of significant differences between the two groups was assessed using a Mann-Whitney test. K. Metagene plots of disome footprints fold-changes for the 5’ TOP and non-5’ TOP transcripts relative to start codons. Alt text: Eleven panels. A and B are metagene line plots of log_2_fold-change against position relative to the start codon and, in a second window, relative to the stop codon, with one coloured line per treatment; A shows monosome signal, B disome signal. C and D present the same data as stacked heatmaps, one horizontal block per treatment, with each row a transcript and colour running from dark blue through teal to yellow on a log2 fold-change scale. E to H each pair a line plot above a heatmap, showing signal split into three transcript clusters: E is UV disome, F UV monosome, G low-dose anisomycin disome and H high-dose anisomycin disome, with cluster sizes of 671, 1342 and 2961 transcripts marked on the heatmap blocks. I is a dot plot of enriched gene ontology terms for cluster 1, with terms listed on the vertical axis against negative log10 FDR on the horizontal axis, dot size encoding gene count and dot colour encoding fold enrichment. J is three pairs of boxplots comparing 85 TOP mRNAs with 85 matched non-TOP mRNAs for RNA-seq counts, Ribo-seq counts and translation efficiency, each pair annotated as not significant. K is three line plots, one per treatment, overlaying TOP and non-TOP disome metagene profiles.

Transcript-level analyses reproduced the positional patterns seen in the metagene plots. Under LD ANS, monosome FC rose sharply downstream of the start codon, peaked at ∼30- 40 nt and declined along the CDS (**Fig. 4A**), and disome FC peaked at ∼60-70 nt (**Fig. 4B**). HD ANS produced the same pattern with maxima closer to the start codon and of lower amplitude. In addition, HD ANS caused a marked loss of monosome signal at the start codon itself, consistent with arrested ribosomes just downstream blocking new initiation events (see model above). UV produced only a modest monosome increase at the start codon and a low-amplitude disome peak, followed by a relative depletion of signal further downstream, consistent with partial ribosome run-off.

The heat maps showed that, under both LD and HD ANS, the post-initiation increase in monosome and disome signal was visible across most transcripts, varying in magnitude but with a strikingly uniform positional pattern (**Fig. 4C, D**). The collision zone is therefore a broadly shared feature rather than the product of a few strongly affected mRNAs. In contrast, the UV-induced disome increase appeared confined to a subset of transcripts (**Fig. 4D**).

To characterise this heterogeneity, we grouped transcripts into three clusters by k-means clustering of their UV disome FC profiles (**Fig. 4E**). Cluster 1 (n = 671) showed a post-initiation disome increase followed by downstream depletion, consistent with collisions accompanied by run-off. Cluster 2 (n = 1,342) showed a broader increase without a discrete peak or downstream depletion, and cluster 3 (n = 2,961) showed little change. UV monosome profiles were similar across the three clusters (**Fig. 4F**), indicating that the heterogeneity is specific to the disome signal. Under LD and HD ANS, all three clusters showed a clear post-initiation disome peak, but its height decreased from cluster 1 to cluster 3 (**Fig. 4G, H**). The transcripts most affected by UV are thus also those most affected by ANS. Under UV, however, only this most responsive subset showed a substantial increase.

Gene ontology analysis of cluster 1 showed strong enrichment for cytoplasmic translation and ribosome biogenesis (**Fig. 4I**), driven largely by mRNAs encoding ribosomal proteins (RPs) and translation factors (**Supplementary Table S2**). Many of these carry 5′ terminal oligopyrimidine (TOP) motifs, which confer mTOR-dependent translational control (50), and they are also among the most abundant and highly translated mRNAs in mESCs (51). To separate these features, we compared TOP mRNAs (n = 85) with a non-TOP set matched for mRNA abundance, ribosome footprint abundance and TE in control cells (**Fig. 4J**). Disome FC profiles were very similar for the two groups under all three conditions (**Fig. 4K**), indicating that collision propensity is associated with high ribosome loading rather than with the TOP motif itself. Consistent with this, post-initiation disome FC increased with TE across the transcriptome for ANS (**Supplementary Fig. S3A-G**).

Together, these analyses indicate that ANS generates a uniform post-initiation collision zone shared across translated mRNAs, with an amplitude that scales with TE. At 30 min after UV, collision accumulation is more heterogeneous and is concentrated on a subset of highly translated transcripts, among which TOP mRNAs are prominent because of their high ribosome loading rather than their TOP motif.

### Site-resolved analyses reveal abundant anisomycin-induced stalling and collision sites but few recurrent UV-dependent sites

The analyses above describe positional patterns and the transcripts affected, but do not resolve individual stalling and collision sites. We therefore identified sites of differential ribosome occupancy transcriptome-wide using an approach adapted from Stein et al. (28), in which occupancy at each codon is compared with that of the remainder of the same CDS between treated and control samples (see Materials and Methods).

Under LD ANS, we detected 1,166 significantly increased ("up") and 142 decreased ("down") monosome sites across 413 genes, and 886 up and 1,240 down disome sites across 408 genes (**Fig. 5A**). Under HD ANS, monosome sites were more numerous (1,354 up, 950 down; 332 genes), whereas disome sites were fewer (658 up, 678 down; 257 genes) (**Fig. 5B**), consistent with earlier arrest and fewer ribosomes progressing into collisions, as predicted by the model above. UV yielded far fewer sites: 101 up and 4 down monosome sites across 87 genes, and 118 up and 56 down disome sites across 96 genes (**Fig. 5C**). Under all conditions, site-containing genes were biased towards abundant, highly translated mRNAs (**Supplementary Fig. S4A, B**; **Supplementary Table S3**), and RP mRNAs were overrepresented among them (**Supplementary Fig. S4C-E**).

**Figure 5.**
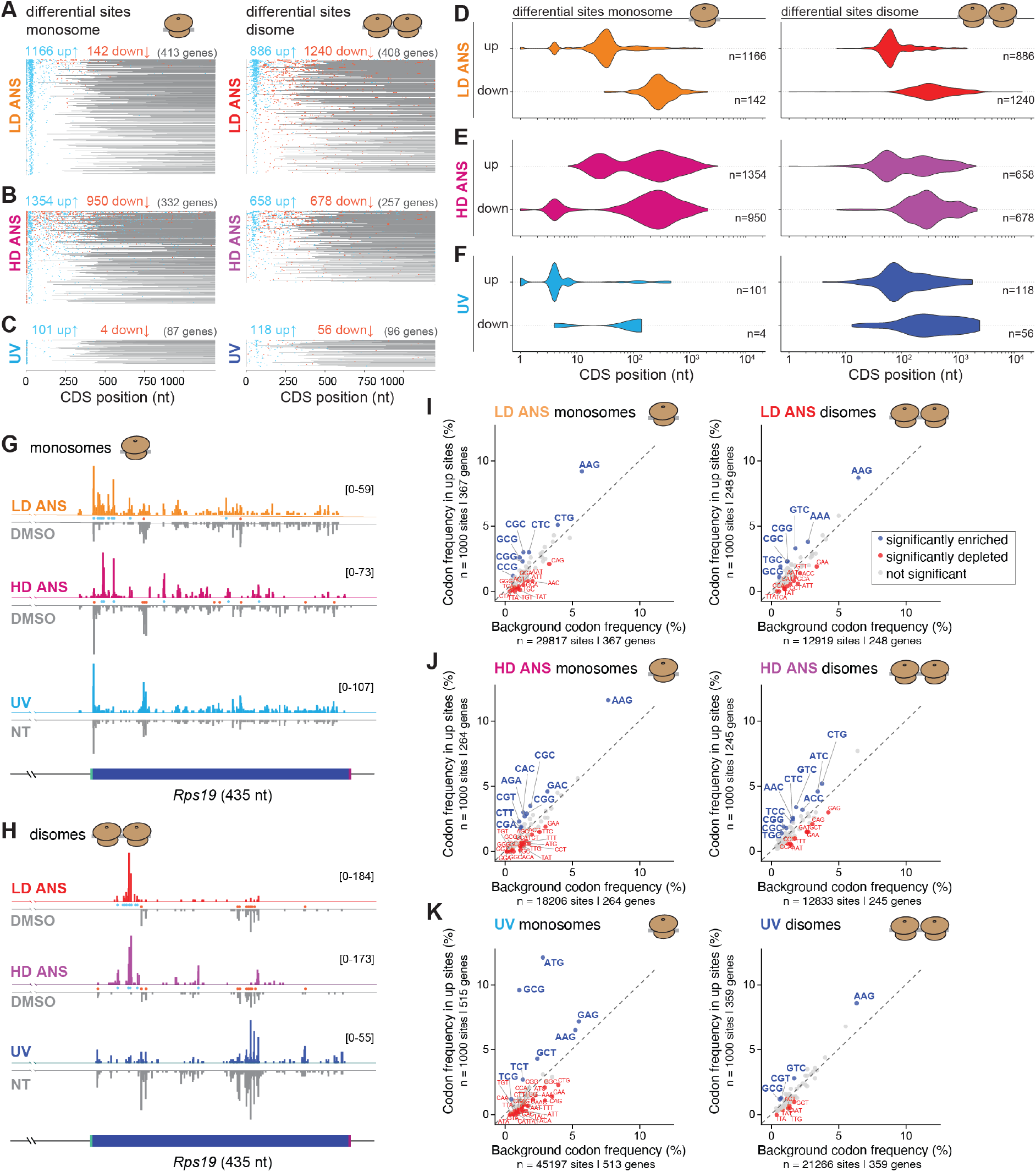
Transcript-wide distribution and sequence features of differential ribosome occupancy sites under ANS and UV conditions. **A.** Distribution of significantly up- and downregulated monosome (left panel) and disome sites (right panel) along individual transcripts under low-dose (LD) ANS conditions. Transcripts are ordered by decreasing number of detected sites, and coding sequences (CDS) are aligned with post-stop-codon regions shown in grey. **B.** As in panel A, for high-dose (HD) ANS treatment. **C.** As in panel A, for UV-treated samples. **D.** Relative positional distribution of significantly up- and downregulated sites along normalized CDS coordinates under LD ANS conditions, monosomes in left panel, disomes in right panel. **E.** As in panel D, for high-dose (HD) ANS treatment. **F.** As in panel D, for UV-treated samples. **G.** Normalised A-site monosome footprint profiles across the *Rps19* transcript under LD ANS, HD ANS, UV, and corresponding control conditions. Replicate signals were pooled for the analysis. Significant sites from the pause score analysis are indicated by blue (up) and red (down) dots between the traces. **H.** As in panel G, showing disome footprint profiles. **I.** Codon composition of A-site positions for the top 1000 most significantly enriched monosome sites (left panel) and disome sites (right panel) under LD ANS conditions compared to background codon frequencies. Significantly enriched (blue) or depleted codons (red) are indicated, using a permutation test. **J.** Codon composition as in panel I, for HD ANS conditions. **K.** Codon composition as in panel I, for UV conditions. Alt text: Eleven panels. A, B and C each show a pair of site maps, monosome on the left and disome on the right, one pair per treatment. Every horizontal grey line is one transcript’s coding sequence plotted against CDS position in nucleotides, with cyan dots marking significantly increased sites and red dots significantly decreased sites; transcript counts and numbers of up and down sites are printed above each map. D, E and F are violin plots of the CDS positions of up and down sites on a logarithmic axis, monosome on the left and disome on the right, one row per treatment, with site numbers to the right of each violin. G and H are genome-browser- style coverage tracks over the *Rps19* coding sequence, 435 nucleotides long, with the treated sample drawn upwards in colour and the control drawn downwards in grey, stacked for the three treatments and with the plotted range given in brackets; G shows monosomes, H disomes. I, J and K are scatter plots of codon frequency in up sites against background codon frequency, monosome on the left and disome on the right, one row per treatment, with a dashed diagonal, codons significantly enriched in blue above the diagonal, significantly depleted codons in red and non-significant codons in grey.

The positions of these sites differed between treatments. Under LD ANS, up-sites were concentrated near the start of coding sequences and down-sites lay further downstream on the same transcripts (**Fig. 5A, D**), as illustrated for both monosome and disome footprints on *Rps19* (**Fig. 5G, H**). This pattern matches the post-initiation collision zone, with relative depletion further downstream. Because each site is tested against the rest of its own CDS, part of this downstream depletion is expected as a consequence of the strong 5′ gain. HD ANS showed the same organisation, with additional up-sites distributed more broadly along coding sequences and a prominent group of monosome down-sites at the start codon (**Fig. 5B, E**), matching the start-codon depletion seen in Fig. 4A. Under UV, most monosome up-sites lay at the start codon, whereas disome up-sites were concentrated within the first ∼100 nt, with down-sites further downstream (**Fig. 5C, F**).

The small number of UV sites contrasts with the biochemical (**Fig. 1F**) and transcript-level (**Fig. 4E**) evidence for UV-induced collisions. Together with the transcript-level heterogeneity, this suggests that UV-induced collisions are distributed over many positions, each occupied too weakly or too variably to reach significance with our approach, rather than concentrated at recurrent hotspots.

To ask whether specific codons are associated with stalling, we analysed the A-site codons (for disomes, that of the leading ribosome) of the 1,000 positions with the highest pause scores in each dataset, including positions below the significance threshold, and compared their codon composition with that of non-significant positions in the same transcripts (**Fig. 5I-K; Supplementary Fig. S5A**). Several codons were enriched in more than one condition, including GC-rich alanine (GCN) and arginine (CGN) codons and the lysine codons AAG and AAA. Some enrichments reflect position rather than codon identity: monosome ATG enrichment under UV corresponds to start-codon dwelling, and GCG enrichment probably reflects the frequent use of GCG near the start of coding sequences (52). Codons enriched specifically under UV (GAC, TCT and TCG in monosomes; CGT in disomes) showed modest effect sizes.

As a complementary approach that does not depend on site selection, we estimated codon-specific dwell times at the E-, P- and A-sites for monosome (**Supplementary Fig. S5B**) and disome data (**Supplementary Fig. S5C**) using Ribo-DT (31), which models footprint counts along coding sequences (see Materials and Methods). Under UV, dwell times corresponded closely to control at all three sites (monosomes, r = 0.85-0.89) and no codon changed significantly, indicating that UV-induced impairment leaves codon-specific elongation kinetics essentially intact. Under ANS this correspondence was lost, but unevenly: in monosome data it was partial at low dose and, at high dose, the E- and P-site landscapes were reorganised to the point of anticorrelation with control, whereas A-site estimates remained positively correlated at both doses. Disome data showed the same contrast between a preserved A-site and poorly corresponding E- and P-sites. This is consistent with an increasing fraction of drug-arrested ribosomes, in which footprint density progressively reports the probability of drug capture rather than native codon kinetics, and indicates that the A site – where anisomycin acts – retains the most interpretable signal.

Within that signal, a coherent codon signature emerged, and it consisted exclusively of reductions. All four glycine codons showed reduced A-site dwell times at both doses and in both monosome and disome data (**Supplementary Fig. S5B, C**), and at the P-site under HD ANS; no codon showed a significant increase in A-site dwell time under either dose. Anisomycin therefore does not generate codon-specific pausing, but acts less efficiently in a particular codon context – a behaviour described for peptidyl transferase centre-targeting antibiotics in bacteria (see Discussion). Changes were also apparent at the E-site, where alanine and serine codons showed increased and arginine and lysine codons decreased dwell times; because these estimates derive from the sites at which correspondence between treated and control was weakest, we regard them as tentative at best. Dwell-time estimates were virtually unchanged when the first 75 nt of coding sequences were excluded (r ≥ 0.996; **Supplementary Fig. S5D**), indicating that the signature does not depend on the post-initiation zone. Notably, the codon enrichments detected among selected pause sites (**Fig. 5I-K**) were not reproduced in this site-selection-free analysis, indicating that they largely reflect where those sites lie within coding sequences rather than the identity of the codon occupying the A-site.

Together, these analyses show that ANS produces numerous, positionally defined stalling and collision sites but no codon-specific pause signature: the only reproducible codon effect is a context in which drug-induced arrest is less efficient. UV-induced impairment at 30 min yields few recurrent sites and no detectable change in codon dwell times. Both observations are consistent with collision position being determined by where ribosomes are when arrest occurs rather than by sequence features of the transcripts.

## Discussion

Low-dose anisomycin is one of the most widely used tools for inducing ribosome collisions and RSR signalling, yet where these collisions form on mRNAs had not been examined systematically. By combining monosome- and disome-resolved ribosome profiling with proteomics of collided-ribosome fractions, we find that ANS-induced collisions are not dispersed along coding sequences but concentrated in a post-initiation zone, and that low- and high-dose ANS follow the same dose-dependent positional rule. At 30 min after UV irradiation, by contrast, collisions are detectable biochemically but appear more dispersed, with few recurrent sites and preferential accumulation on highly translated mRNAs.

The current view holds that low ANS concentrations lead to sparse ribosome stalls along transcripts, against which upstream ribosomes collide, whereas high concentrations freeze ribosomes in place and prevent collisions (9,10). Our data support a different picture. Under both regimes, stalled monosomes and disomes accumulate within roughly the first 20 codons, visible in metagene representations, on individual transcripts and at discrete sites. Higher doses shift this zone towards the start codon and reduce its amplitude without abolishing it. Consistent with this, high concentrations of elongation inhibitors had previously been inferred to arrest ribosomes shortly after start-codon recognition, on the basis that they elevate RNF10- dependent 40S ubiquitylation (47); the profiling data place that shift directly.

We propose that this pattern arises from the interplay between a dose-dependent probability of drug-induced arrest at each elongation cycle and continued initiation. Ribosomes arrested within about one footprint length of the start codon occlude the initiation site and remain as monosomes. Those that progress further before arresting allow a new ribosome to initiate and collide behind them (**Fig. 6A**). According to this view, the dose of ANS determines how far ribosomes advance before arrest, and initiation determines the supply of trailing ribosomes (**Fig. 6B**). Independent support comes from the dwell-time analysis: in monosome data, the correspondence between treated and control codon dwell times declined progressively from UV to low-dose and then high-dose ANS, as expected if an increasing fraction of footprints reports drug capture rather than native elongation kinetics. The model makes testable predictions, for instance on how the zone should respond to intermediate doses, longer exposures or reduced initiation, that future experiments can address.

**Figure 6.**
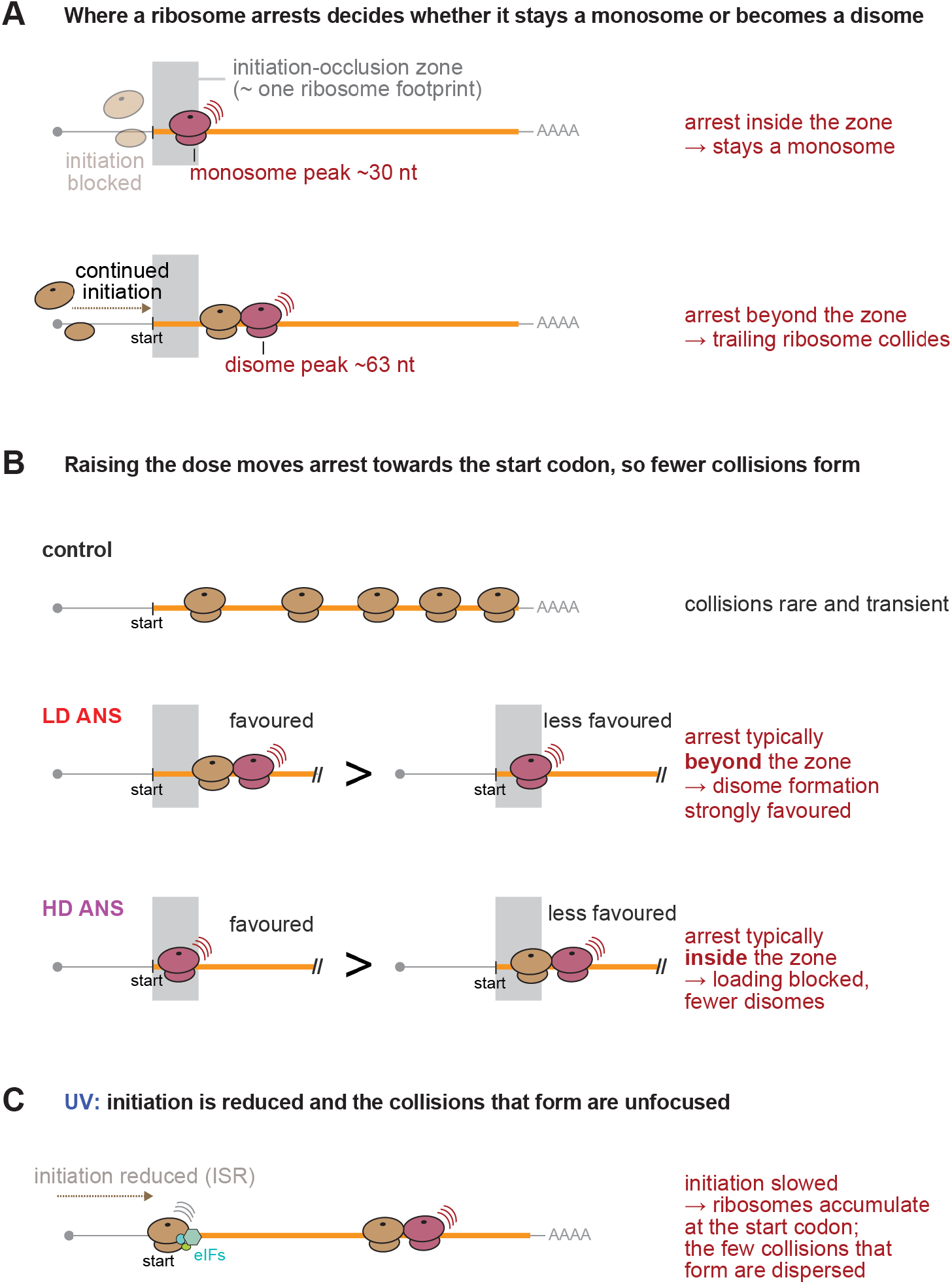
A dose-dependent positional rule for anisomycin-induced ribosome collisions, and the contrasting UV response. **A.** Schematic of the proposed rule. Ribosomes arrest stochastically during elongation at a rate set by the ANS concentration. A ribosome arrested within approximately one footprint of the start codon (grey band, "initiation-occlusion zone") continues to occlude the initiation site, prevents a further ribosome from loading (pale ribosomal subunits, "initiation blocked") and is recovered as a monosome. A ribosome that advances beyond this zone before arresting vacates the initiation site, allowing a new ribosome to load and run into it, and is recovered as a disome. The two fates account for the different positions of the monosome and disome peaks: ∼30 and ∼63 nt downstream of the start codon under LD ANS, and ∼24 and ∼48 nt under HD ANS (Fig. 3E, J); peak positions are approximate (see Discussion). Throughout the figure, tan ribosomes are elongating, dark red ribosomes are arrested, red radiating lines denote arrest and collision, and all drawings are schematic and not to scale. **B.** Schematic of the ANS dose dependence. Under control conditions collisions are rare and transient. Under LD ANS, arrest beyond the occlusion zone is favoured over arrest within it, so a trailing ribosome frequently loads and collides. Under HD ANS the same two outcomes occur with the weighting reversed: arrest typically falls within the zone, further loading is blocked and fewer disomes form – consistent with the reduced but readily detectable disome signal (Fig. 1E**, 3H**) and with the loss of monosome occupancy at the start codon (Fig. 4A**, 5B**). "Favoured" and "less favoured" denote relative likelihoods under the model, not measured ratios. Double slashes mark truncated transcripts. **C.** Schematic of the UV response at 30 min. ISR activation (Fig. 1G) reduces initiation, and ribosomes accumulate at the start codon (Fig. 3K**, 5C**) in association with initiation factors (Fig. 2E); whether these complexes are 80S ribosomes retaining initiation factors or 40S- containing initiation intermediates was not resolved. Fewer ribosomes load onto coding sequences, and the collisions that do form (Figures 1F**, 4D**) are infrequent and not positionally focused. The nature of the impediments at which UV-induced collisions form was not determined here. Alt text: Three schematic panels, each with a title above it. A shows two mRNAs drawn as horizontal lines with a start codon, a poly-A tail and a grey vertical band marking an initiation- occlusion zone about one ribosome footprint wide. On the upper mRNA a dark red arrested ribosome sits inside the band, pale ribosomal subunits to its left indicate blocked initiation, and the monosome peak is labelled at about thirty nucleotides. On the lower mRNA the arrested ribosome sits beyond the band, a tan ribosome has loaded behind it, and the disome peak is labelled at about sixty-three nucleotides. B shows the same mRNA under three conditions: a control with five evenly spaced tan ribosomes, then low-dose and high-dose anisomycin, each drawn as a favoured and a less favoured outcome separated by a greater-than sign, with arrest occurring beyond the grey band under low dose and inside it under high dose. C shows a single mRNA after UV with a reduced-initiation arrow, a ribosome carrying small teal initiation factors stalled at the start codon, and one collided ribosome pair further along. Throughout, tan ribosomes are elongating, dark red ribosomes are arrested and red radiating lines mark arrest and collision; annotations in red to the right of each panel state the outcome.

Two practical consequences follow from these findings. First, HD ANS should not be regarded as collision-free. It produces fewer collisions, but with the same organisation and with detectable JNK phosphorylation, in contrast to observations in other cell systems (10). Second, because LD ANS-induced collisions are concentrated close to the start codon, they occur where initiation factors are still present on the ribosome (46) and where collisions are well placed to engage feedback on initiation (32,45). Studies using this paradigm therefore examine a collision population with specific positional properties, which should be considered when generalising from it to collisions arising elsewhere in coding sequences.

Superimposed on this dominant positional pattern, ANS also acted in a codon-dependent manner, but not in the way a sequence-specific stalling signal would. The only reproducible codon effect was a reduction: all four glycine codons showed shorter A-site dwell times at both doses and in both monosome and disome data, and no codon showed a significant increase. ANS therefore does not create codon-specific arrest sites; rather, there is a codon context in which drug-induced arrest is less efficient. This parallels the context specificity described for peptidyl transferase centre-targeting antibiotics in bacteria (53,54), which likewise act unevenly along a transcript without generating defined arrest positions. E-site preferences have also recently been reported for ANS (55); our data are compatible with this, although the E-site estimates are the least well determined in our analysis. Consistent with the absence of a positive codon signature, the enrichments we detected among selected pause sites were not reproduced once site selection was removed, indicating that they largely reflect where those sites lie within coding sequences rather than the identity of the codon in the A-site. Codon context and positional accumulation are thus separable – the dwell-time signature was virtually unchanged whether or not the post-initiation zone was included – and where ANS- arrested ribosomes come to rest is determined principally by when arrest occurs during elongation, not by the sequence at which it occurs.

The UV response differed in kind, not only in degree. Thirty minutes after irradiation the dominant signature was at initiation: eIF2α phosphorylation (**Fig. 1G**), accumulation of ribosomes at the start codon, co-sedimentation of initiation factors with monosome and disome fractions, and no detectable codon-specific change in elongation dwell time. Blocking progression of scanning or elongating ribosomes past the start codon marks 40S subunits for iRQC (47), and ISR activation is itself a trigger; the ribosome population we observe accumulating at the start codon after UV is therefore a candidate iRQC substrate, although we did not assay ribosomal ubiquitylation. Collisions did form as well – nuclease-resistant disomes and trisomes were readily detectable – but they were few, not positionally focused, and concentrated on a subset of transcripts (**Fig. 6C**). Because ISR activation is expected to reduce initiation, and hence the supply of trailing ribosomes, collisions should be infrequent wherever impediments arise; the same two determinants that shape the ANS landscape therefore account for the UV one, with initiation rather than arrest position as the throttled variable. The nature of those impediments is not addressed by our data. Stochastically distributed RNA photolesions would be consistent with the dispersed, non-recurrent pattern we observe (12,13), but we have not measured them, and photolesions (11) may additionally affect library preparation. Our analysis also captures a single time point, so the landscape may change with recovery. We therefore regard the UV landscape as descriptive of the 30 min time point rather than as evidence that UV-induced collisions lack positional organisation.

Under both ANS and UV, collision signal scaled with translation efficiency, and the transcripts most affected by UV were also those most affected by ANS. TOP mRNAs were prominent among them, but matched non-TOP transcripts behaved similarly, suggesting that ribosome loading rather than the TOP motif determines collision propensity. Highly translated mRNAs may thus form a preferred substrate pool for collisions, as previously suggested (6,56). Highly covered transcripts are, however, also more readily detected in transcript- and site-level analyses, and separating biological susceptibility from detection sensitivity will require approaches that control for coverage.

We found EDF1 enriched in disome fractions under all conditions, consistent with its role as a general collision sensor (35), whereas ZNF598 and RQC execution factors were most clearly enriched after UV, and components of ER-associated RQC and UFMylation (40) co-sedimented with impaired ribosomes under LD ANS and UV. Initiation factors, notably eIF3, co-sedimented with monosome and disome fractions, most prominently after UV. This is consistent with impairment shortly after initiation, but under UV it may also reflect 40S- containing initiation intermediates accumulating as a result of ISR activation. Distinguishing these possibilities will require approaches that resolve subunit composition directly.

Our approach has three main limitations that we wish to put into perspective in the following. First, ribosome states were preserved with elongation inhibitors – cycloheximide for ribosome profiling, ANS for gradient analyses. Cycloheximide can generate 5′-proximal artefacts (57,58), but controls processed identically showed no post-initiation accumulation, so the collision zone is ANS-dependent. Nevertheless, we cannot exclude that the two strategies capture partly different populations of collided ribosomes. Second, footprints were selected within fixed length windows and A-site positions inferred from fixed offsets, and anisomycin alters how far nucleases trim footprints (59). The post-initiation zone, its presence in both monosome and disome data, and its absence under UV are robust to these choices; the peak positions and the dose-dependent shift are approximate. Third, our measurements are relative rather than absolute – profiling signals are library-normalised, the proteomic data report co-sedimentation rather than direct binding, and both site detection and dwell-time estimation scale with coverage – so magnitudes should not be compared quantitatively between conditions.

Two questions remain open. ZAKα dependence was not tested in mESCs, although ZAKα mediates JNK activation under ANS and UV in other systems (7,8,10,12). And these landscapes describe one cell type, one ANS exposure and a single time point after UV; how they evolve with exposure, recovery and cell type remains to be established.

In summary, low-dose anisomycin does not scatter collisions along transcripts but confines them to a post-initiation zone whose position is set by the dose, and high-dose anisomycin follows the same rule rather than abolishing collisions. UV acts not on where ribosomes arrest but on how many follow behind them: by limiting initiation it reduces the supply of trailing ribosomes, so collisions are few and without positional focus. Where collisions form, and not only how many, thus shapes what these paradigms report, and will matter for understanding how collisions are converted into stress signals.

## Data Availability

Mass-spec data is available on ProteomeXchange (https://www.ebi.ac.uk/pride/login with Project accession: PXD077189, token **********). Sequencing data is available on NCBI GEO Omnibus (GSE344663, token **********, for monosome/disome Ribo-seq data; GSE344668, token **********, for RNA-seq data).

Computational code is available at: https://github.com/gatfieldlab/Snakemake_pipeline for the mapping of sequencing data; https://github.com/gatfieldlab/ANS_UV/ for downstream analyses on the mapped data; and https://github.com/cgob/codonDT_snakemake for the RiboDT analyses.

## Supplementary Data statement

Supplementary Data are available at *NAR* Online.

## Acknowledgements

We would like to thank the University of Lausanne Proteomics Analysis Facility and the

Genomic Technologies Facility for expert advice and services that were essential to this work. We also thank members of the Gatfield lab for comments on the manuscript.

## Author Contributions Statement

SC – Writing original draft, Visualization, Validation, Conceptualization, Formal analysis, Investigation, Methodology. JS – Writing review and editing, Visualization, Validation, Formal analysis, Investigation, Methodology. VR – Writing review and editing, Investigation, Formal analysis, Methodology. FN – Supervision. CG – Writing review and editing, Visualization, Formal analysis, investigation. DG **–** Writing original draft, Writing review and editing, Visualization, Funding acquisition, Conceptualization, Formal analysis, Supervision, Project administration.

