## Supplemental Figures for "Anisomycin-induced ribosome collisions follow a dose-dependent positional rule, distinct from the UV-induced collision landscape"

\* corresponding author

Figure S1

A

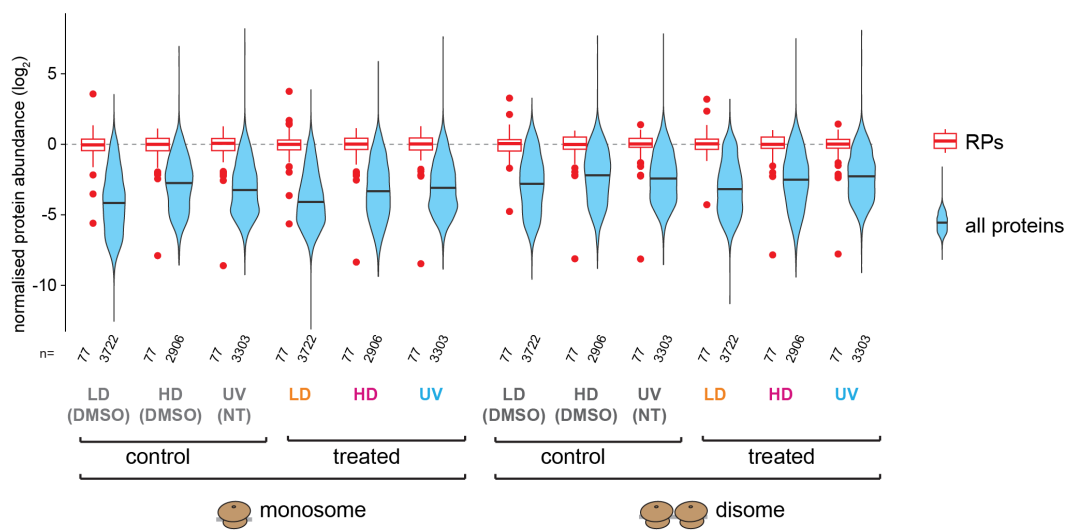

B

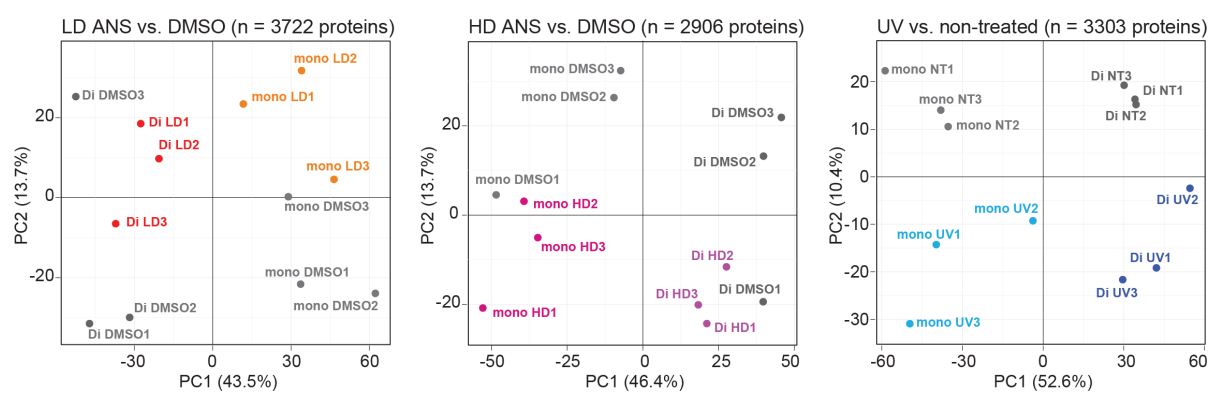

**Figure S1. Quality control and normalisation of ribosome-associated proteomics datasets**

**A.** Distribution of ribosomal protein (RP)-normalised protein abundances across monosome and disome fractions under control, LD ANS, HD ANS, and UV conditions. The median RP abundance was set to 0 on the $\log_2$-scale and the whole proteome adjusted accordingly. While this normalisation procedure was applied to each sample individually, the graph here shows the means over triplicates per condition. The normalised protein amounts were used to calculate fold-changes and significance (t-test) for the data shown in Figure 2 and Table S1.

**B.** Principal component analysis (PCA) of proteomic datasets. Samples are coloured according to treatment condition. Fraction identity (mono/di) dominates PC1, with treatment separating on PC2 for the UV comparison; replicate clustering was less clean for the ANS comparisons.

Alt text: Two panels. A plots normalised protein abundance on a $\log_2$-scale for 12 sample groups along the horizontal axis: control and treated monosome fractions for low-dose anisomycin, high-dose anisomycin and UV, followed by the same six groups for disome fractions. Each group is drawn as a blue violin showing all proteins with a narrow red boxplot of ribosomal proteins overlaid, and the number of proteins in each is printed beneath. B is three principal component analysis scatter plots, one per comparison, plotting the first against the second principal component with the percentage of variance explained on each axis; individual monosome and disome samples are plotted as labelled points, treated samples in colour and controls in grey.

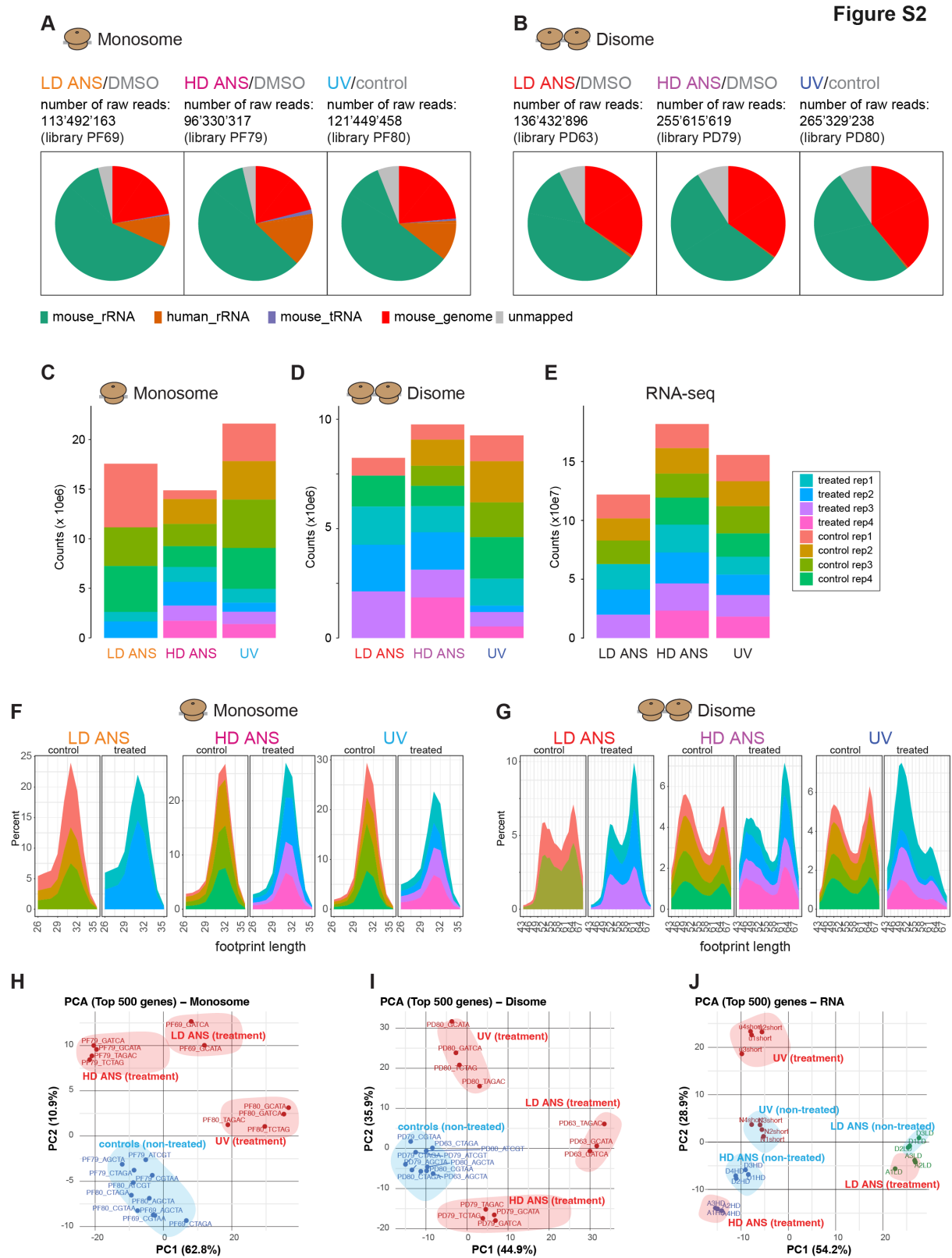

**Figure S2. Sequencing quality metrics and overview of ribosome profiling datasets.**

**A.** Numbers of overall raw reads obtained for monosome ribosome profiling libraries for each independent condition, and outcome in terms of mouse\_genome reads that formed the input for subsequent analyses.

**B.** As in panel A, for disome ribosome profiling libraries.

**C.** Number of monosome footprint reads – after de-duplication, removal of multimapping reads etc. – that are then used for further analyses in the study.

**D.** As in panel C, for disome ribosome profiling libraries.

**E.** Number of RNA-seq reads per condition/replicate used for the analyses of the study.

**F.** Footprint length distribution for monosome reads used in the analyses.

**G.** As in panel F, for disome data.

**H.** Principal component analysis for monosome data.

**I.** Principal component analysis for disome data.

**J.** Principal component analysis for RNA-seq data.

Alt text: Ten panels. A and B are two rows of three pie charts showing the fate of sequencing reads for monosome and disome libraries respectively, one chart per treatment, segmented into mouse ribosomal RNA, human ribosomal RNA, mouse transfer RNA, reads mapping to the mouse genome and unmapped reads, with total raw read counts and library identifiers printed above. C, D and E are stacked bar charts of read counts for monosome, disome and RNA-seq libraries, with one bar per treatment subdivided into four treated and four control replicates. F and G are footprint length distributions plotted as filled area curves of percentage against footprint length, with control and treated panels side by side for each treatment; F covers monosome lengths from 26 to 35 nucleotides and shows a single peak near 30, G covers disome lengths and shows two peaks. H, I and J are principal component analysis scatter plots for monosome, disome and RNA samples, plotting the first two components with individual libraries labelled and treatment and control groups enclosed in shaded ellipses.

**Figure S3**

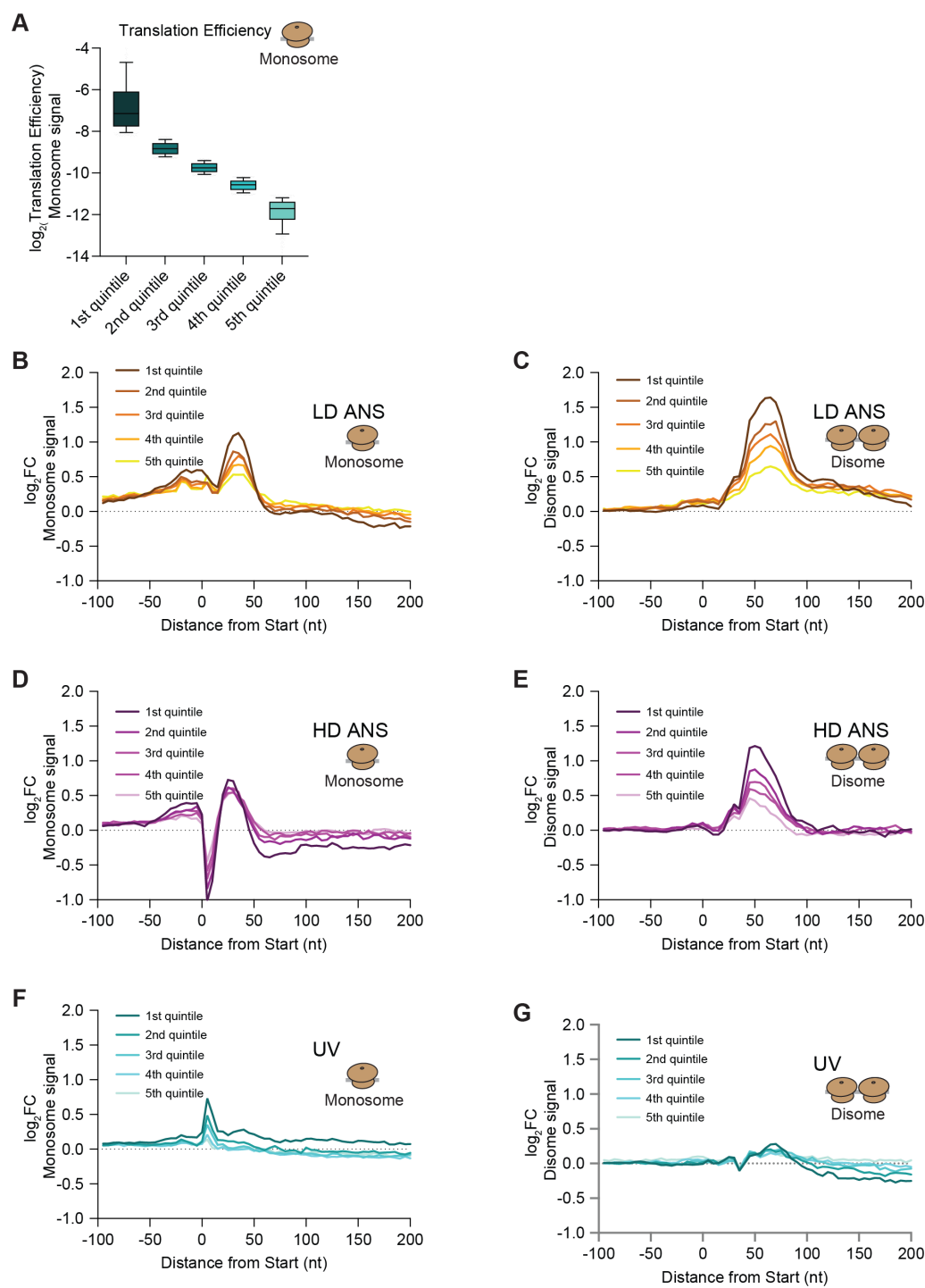

**Figure S3. Translation efficiency correlation with fold-change monosome and disome signals.**

**A.** Boxplots showing the distribution of transcripts across quintiles ranked by steady-state translational efficiency (TE), determined in the control sample of the low-dose anisomycin experiment.

**B.** Metagene plots showing fold changes in monosome footprints across transcripts grouped by steady-state TE quintile under low-dose anisomycin condition.

**C.** As in panel B, for disome footprints.

**D.** As in panel B, under high-dose anisomycin condition.

**E.** As in panel C, under high-dose anisomycin condition.

**F.** As in panel B, under UV-treated condition.

**G.** As in panel C, under UV-treated condition.

Alt text: Seven panels. A is a series of five boxplots of log<sub>2</sub> translation efficiency for transcripts grouped into quintiles, shaded from dark to light teal and decreasing stepwise from left to right. B to G are metagene line plots of log<sub>2</sub> fold-change against distance from the start codon, from minus one hundred to plus two hundred nucleotides, each containing five lines coloured by translation efficiency quintile. B and C show monosome and disome signal under low-dose anisomycin, D and E under high-dose anisomycin, and F and G under UV; a dotted horizontal line marks zero in every panel.

Figure S4

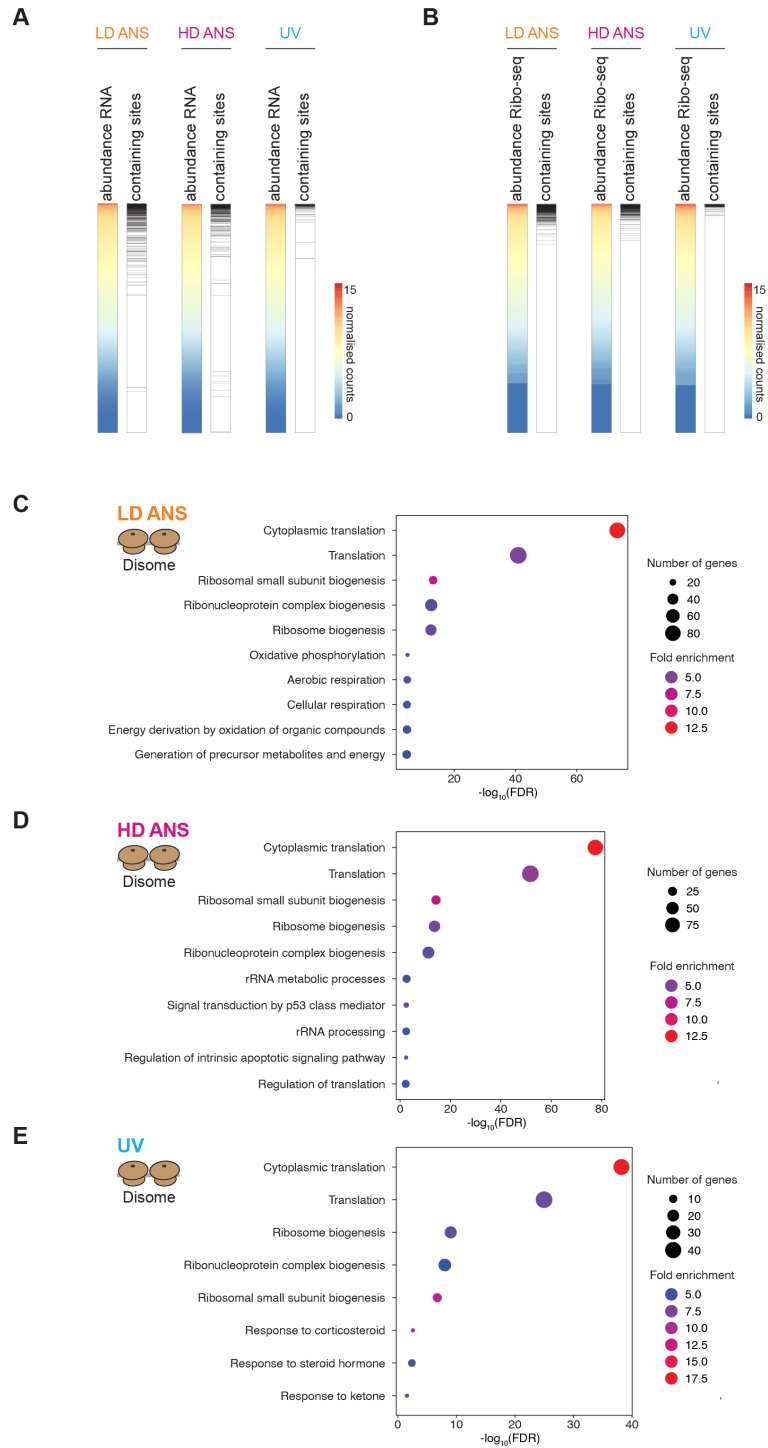

**Figure S4. Analysis of site-containing transcripts**

**A.** Relationship between expression levels (from RNA data) and significant monosome stalling sites across all transcripts. Heatmaps on the left are sorted by expression level in treated conditions; to the right of each heatmap, the panels mark by black lines the transcripts containing monosome significant stalling sites.

**B.** As in panel A, but using expression levels from Ribo-seq data.

**C.** Gene ontology enrichment analysis of transcripts containing significant disome sites under LD ANS conditions.

**D.** Gene ontology enrichment analysis of transcripts containing significant disome sites under HD ANS conditions.

**E.** Gene ontology enrichment analysis of transcripts containing significant disome sites under UV conditions.

Alt text: Five panels. A and B each show three pairs of narrow vertical strips, one pair per treatment. The left strip of each pair is a heatmap of transcript abundance, coloured from blue at low counts through yellow to red at high counts, with transcripts ordered from most to least abundant; the right strip is a black-and-white rug in which each black line marks a transcript containing a differential site. A ranks transcripts by RNA abundance, B by Ribo-seq abundance. C, D and E are dot plots of enriched gene ontology terms for disome site-containing transcripts under low-dose anisomycin, high-dose anisomycin and UV respectively, with terms listed on the vertical axis against negative $\log_{10}$FDR on the horizontal axis, dot size encoding the number of genes and dot colour encoding fold enrichment.

Figure S5

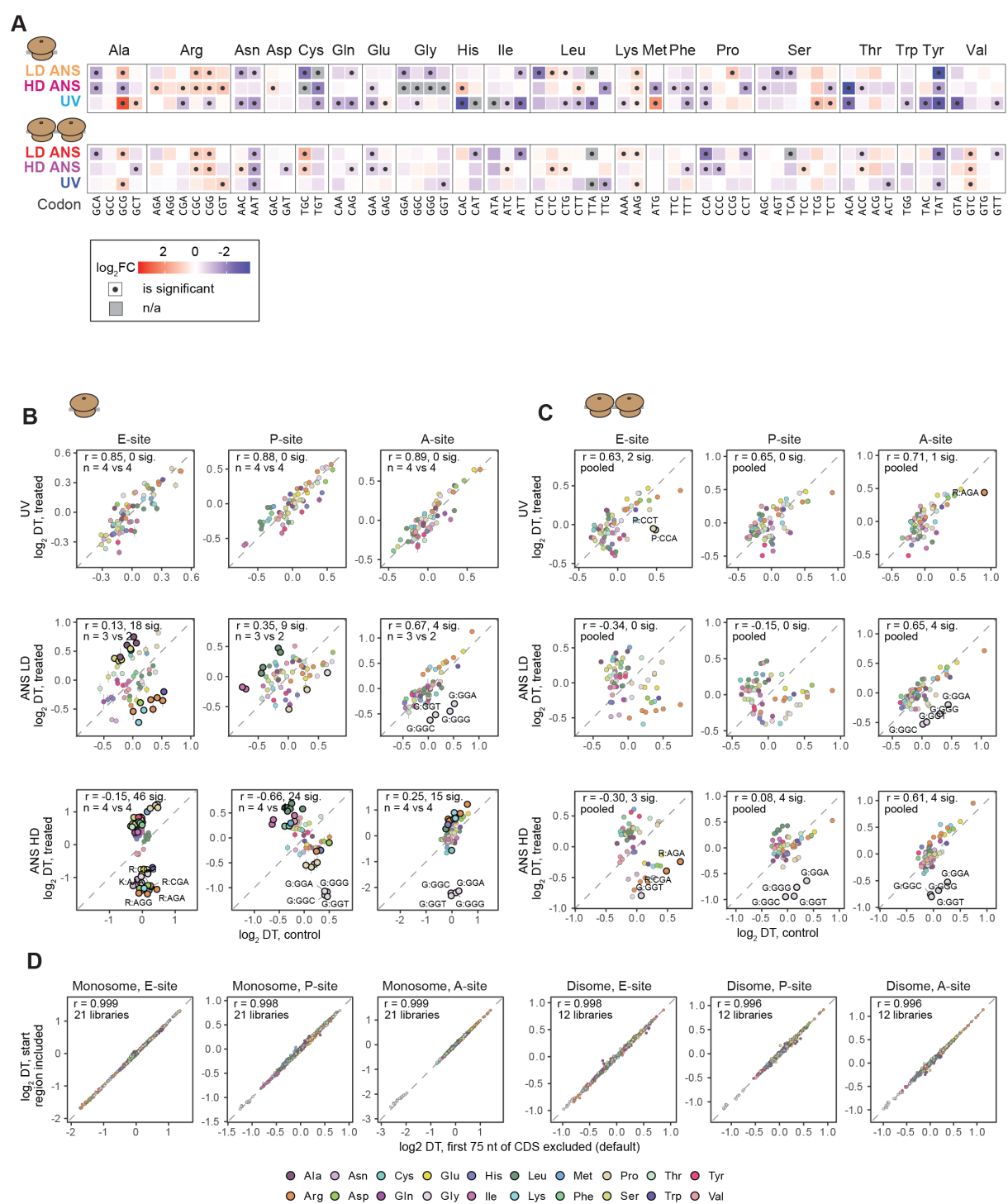

**Figure S5. Codon analysis of monosome and disome footprint data**

**A.** Heatmap representation of enrichments shown in Figure 5 I-K, indicating the log<sub>2</sub>FC on the indicated colour scale, with a dot in the box specifying statistical significance calculated using permutations test.

**B.** Codon dwell times (log<sub>2</sub>) at the E-, P- and A-sites, estimated with Ribo-DT using a single-codon model that includes the post-initiation zone, using monosome data. Treated vs. control for UV, low-dose ANS and high-dose ANS. Points are coloured by amino acid; the dashed line is the identity; $r$, Pearson correlation. Mean ± SEM of biological replicates fitted separately ($n$, control vs treated). Outlined codons differ significantly (t-test, Benjamini-Hochberg-adjusted $p < 0.05$, change $> 0.5 \log_2$); labelled codons are also outliers (modified z-score $> 2.5$).

**C.** Similar to panel B, but for Disomes with replicates pooled per condition. Labelled codons differ significantly (Wald test, Benjamini-Hochberg-adjusted $p < 0.05$) and are outliers (modified z-score $> 3$).

**D.** Dwell times with the first 75 nt of coding sequences excluded (x) or included (y), for each library and codon (monosomes, 21 libraries; disomes, 12 libraries).

Alt text: Four panels. A is a pair of heatmap blocks, monosome above and disome below, each with three rows for low-dose anisomycin, high-dose anisomycin and UV and sixty-one columns, one per sense codon, grouped and bracketed by amino acid from alanine to valine; cells are shaded on a red-to-blue $\log_2$ fold-change scale, a dot marks significance and grey marks codons with no estimate. B and C are three-by-three grids of scatter plots for monosome and disome data respectively, with rows for UV, low-dose anisomycin and high-dose anisomycin and columns for the E, P and A sites; each plots $\log_2$dwell time in treated against control samples with a dashed diagonal, points coloured by amino acid, and the correlation coefficient, number of significant codons and replicate numbers annotated in the corner. D is six scatter plots comparing $\log_2$dwell times estimated with the start region included against the default in which the first seventy-five nucleotides of the coding sequence are excluded, for the E, P and A sites of monosome and disome data, each annotated with a correlation coefficient and the number of libraries. An amino acid colour key runs along the bottom of the figure.
